# Mesenchyme modulates three-dimensional, collective cell migration of liver cells *in vitro* - a role for TGFß pathway

**DOI:** 10.1101/2020.09.27.315580

**Authors:** Ogechi Ogoke, Osama Yousef, Cortney Ott, Allison Kalinousky, Lin Wayne, Claire Shamul, Shatoni Ross, Natesh Parashurama

**Author notes:** Correspondence Author: Natesh Parashurama, 906 Furnas Hall, Buffalo, NY 14260.

## Abstract

Three dimensional (3D) collective cell migration (CCM) is critical for improving liver cell therapies, eliciting mechanisms of liver disease, and modeling human liver development/ organogenesis. Here, we modeled liver organogenesis to induce 3D CCM and improve existing models. The liver diverticulum, normally surrounded by septum transversum mesenchyme (STM) at E8.5, was modeled with a miniature liver spheroid surrounded by mesenchymal cells and matrix. In mixed spheroid models with both liver and uniquely MRC5 (fetal lung) fibroblasts, we observed co-migration of cells, and a significant increase in length and number of liver spheroid protrusions, and this was highly sensitive to TGFB1 stimulation. To understand paracrine effects between MRC-5 cells and liver, we performed conditioned medium (M-CM) experiments. Interestingly, the addition of M-CM increased liver 3D CCM, with thin, 3D, dose-dependent branching morphogenesis, an upregulation of Twist1, and a sensitivity to a broad TGFB inhibitor. To test the effects of cell-cell interactions of 3D CCM, the STM was modeled with a spheroid of MRC-5 cells, and we performed co-spheroid culture of liver with MRC-5. We observed a complex morphogenesis, whereby thin, linear, 3D liver cell strands attach to the MRC-5 spheroid, anchor, and thicken to form permanent and thick anchoring contacts between the two spheroids. We also observed spheroid fusion, a form of interstitial migration. In conclusion, we present several novel cultivation systems that induce distinct features of 3D CCM, as judged by the presence of branching, linearity, thickness, and interstitial migration. These methodologies will greatly improve our molecular, cellular, and tissue-scale understanding of liver organogenesis, liver diseases, and liver cell therapy, and will serve as a tool to bridge conventional 2D studies and preclinical *in vivo* studies.

## INTRODUCTION

Liver (epithelial) cell migration plays important roles in liver organogenesis, disease, and therapy. During liver organogenesis (E8.5), hepatic endoderm lining the liver diverticulum undergoes epithelial to mesenchymal transition (EMT), and initiates three-dimensional (3D) collective cell migration CCM), forming hepatic cords, and migrating into the surrounding connective tissue termed the septum transversum mesenchyme (STM) to form the liver bud (Bort, Signore et al. 2006). Hepatic cords are required for liver organogenesis. Studies demonstrate that hepatic endoderm co-migrates with endothelial cells (Matsumoto, Yoshitomi et al. 2001), and STM cells, undergoes branching suggestive of early 3D branching morphogenesis (Watt, Zhao et al. 2007), and eventually forms primitive cell sheets consisting of hepatoblasts (Sosa-Pineda, Wigle et al. 2000). At later stages of liver organogenesis (E17), recent studies have generally highlighted fetal hepatoblasts proliferation (Boylan, Francois-Vaughan et al. 2017). However, studies of rat fetal hepatoblast expression indicate that genes associated with 3D CCM, morphogenesis, and extracellular matrix remodeling, are highly upregulated in fetal hepatoblasts, including lysyl oxidase, lysosomal mannosidase beta A, tenascin (Tnc), semaphorin IV isoform, procollagen C-endopeptidase enhancer 2, adam19, glypican-3, and cathepsin L (Petkov, Kim et al. 2000). This strongly suggests that interstitial migration plays a role in fetal liver growth. An example of liver disease in which 3D CCM is critical is in hepatocellular carcinoma (HCC), and studies demonstrate the importance of local spread and metastasis, including worsened prognosis and increased treatment resistance (Yang, Chen et al. 2009). Finally, liver 3D CCM is important in the setting of liver cell therapy, where either adult or fetal hepatocytes are injected into acute and chronic liver disease models to repopulate the diseased liver. In this case, non-invasive imaging has demonstrated that transplanted hepatocytes enter large veins and then capillaries within hours, migrate across the liver sinusoids, and through liver tissue (Koenig, Stoesser et al. 2005); (Rajvanshi, Kerr et al. 1996); (Gupta, Rajvanshi et al. 1999). Thus, since there are several phases of liver 3D CCM, including co-migration, branching, and interstitial migration, there is a clear need for further studies that identify mechanisms in human cells.

Although molecular mechanisms that drive 2D vs. 3D liver cell migration have not been elucidated, many studies have elicited molecular pathways that drive migration. In the liver diverticulum, 3D CCM occurs in part in response to secreted FGF2 from the cardiac mesoderm, BMP4 from the STM, and endothelial cell interactions (Ledouarin 1964);(Gualdi, Bossard et al. 1996);(Rossi, Dunn et al. 2001);(Matsumoto, Yoshitomi et al. 2001), and migration-associated transcription factors include Hex (Martinez Barbera, Clements et al. 2000), Prox1 (Sosa-Pineda, Wigle et al. 2000), and Tbx3 (Suzuki, Sekiya et al. 2008). For example, GATA4 -/- embryos do not form the STM, despite the presence of endothelial cells, and this leads to a lack of hepatic cord formation and subsequent liver development (Watt, Zhao et al. 2007);(Delgado, Carrasco et al. 2014). Along these lines. conditional knockout of Hlx disrupts liver formation, and was determined to be a strong mediator of necessary mesenchymal-epithelial cell interactions (Hentsch, Lyons et al. 1996). Moreover, in HGF -/- mutants without HGF secretion, presumably by the STM, the liver was hypoplastic, exhibited apoptosis, and decreased hematopoiesis (Uehara, Minowa et al. 1995);(Schmidt, Bladt et al. 1995). Further, in Smad2/3 (+/-) double heterozygotes, the liver is hypoplastic and anemic, and the phenotype demonstrates hepatoblast clustering, and a loss of and mis-localization of beta-1 integrin expression (Weinstein, Monga et al. 2001). Thus, genetic studies collectively indicate reciprocal, molecular interactions between hepatoblast and mesenchyme are necessary for liver growth. In contrast to these studies of 3D CCM in liver organogenesis, current *in vitro* studies of human hepatic migration are limited mostly to 2D assays of highly migratory HCC cells combined with *in vivo* heterotopic or orthotopic tumor models, but typically don’t include supporting (mesenchymal) cell types. The classic assays include wound healing (scratch) assays, as well as transwell assays. Transwell assays can be considered a 3D single cell migration assay, but don’t account for all the types of CCM that occur in biology. Using this approach, scientists have demonstrated various factors that control 2D HCC migration including TGFß1 (Fransvea, Angelotti et al. 2008);(Ng, Tung-Ping Poon et al. 2013), c-Myc (Zhao, Jian et al. 2013), YAP (Fitamant, Kottakis et al. 2015), goosecoid (Xue, Ge et al. 2014), actopaxin (Biname, Lassus et al. 2008), and more recently, miRNAs (miRNA-135a (Zeng, Liang et al. 2016), miRNA-338-3p (Chen, Liang et al. 2017), miRNA-1301(Yang, Xu et al. 2017), mir-665 (Hu, Yang et al. 2018)). Thus, 3D murine migration during liver organogenesis, and 2D modeling in HCC have thus far provided potential molecular mechanisms driving or inhibiting liver 3D CCM.

Despite this progress, it is unclear if these mechanisms would confer similar phenotypes in human 3D CCM systems. This is because fundamental mechanisms of 2D and 3D collective CCM can be distinguished, and 3D CCM has a more varied collection of migration modes, modes (Friedl and Gilmour 2009);(Wu, Giri et al. 2014). The modeling of mesenchyme in liver epithelial/mesenchymal interactions for 3D CCM is in its infancy. In other epithelial (non-liver) systems, studies of non-transformed mammary cancer cells demonstrated that downregulation of Tpm2.1 led to blocked 3D CCM, but paradoxically, enhanced single cell migration (Shin, Kim et al. 2017). Spheroid studies of human breast and head and neck cancer also demonstrate increased diversity of modes of 3D CCM when changing physiological conditions, suggesting the importance of modeling 3D mechanisms (Lehmann, Te Boekhorst et al. 2017). 3D epithelial-mesenchymal interactions, which are critical in the case of the liver, have recently been modeled employing mixed spheroid models. These mixed-cell spheroids demonstrate 3D CCM and co-migration of hepatic (Huh-7) with Lx-2 (mesenchymal) cells in the presence of MG (MG), and similarity to the profibrotic environment in liver cancer (Khawar, Park et al. 2018), and enabled an improved understanding of chemo-resistance. Finally, human stem cell-derived hepatic endoderm has been used in 3D modeling the human liver bud. Importantly, these studies support that 3D co-culture hepatic endoderm with mesenchyme results in a completely different trajectory of gene expression involving epithelial migration genes and epithelial/ mesenchymal interactions (Camp, Sekine et al. 2017). These isolated studies support the idea that 3D models of liver migration are an important part of the toolbox and that mesenchyme-modeling is critical.

Despite a fundamental role of 3D CCM in liver organogenesis, disease, and liver cell therapy, and the fact that 3D migration mechanisms are different than 2D mechanisms, there remains a lack of model systems for liver 3D CCM. Models of human liver cell migration are important for understanding and accurately modeling liver 3D CCM, and existing systems have their limitations. To understand the process more fundamentally, we engineered several different systems that exhibit distinctive characteristics and morphogenetic features of liver 3D CCM, that were distinguished in terms of the length, thickness, and kinetics of 3D CCM. Further, our models demonstrate distinct aspects of 3D CCM and liver morphogenesis, including co-migration, linear motion, branching, and interstitial migration. When testing different mesenchymal cell types, we more thoroughly define the role for MRC5 (lung fetal) fibroblasts in inducing liver 3D CCM. In this model, we identify that TGFß pathway plays a mechanistic role in liver CCD migration in MRC5-induced fibroblasts. To further model interstitial aspects of migration, we develop a novel spheroid co-culture system which demonstrate the range of CCM that occurs between these cell types. These systems enable robust modeling of 3D CCM which will improve our molecular and cellular understanding of liver organogenesis, cancer, and therapy.

## METHODS

### Reagents/materials

DMEM (Thermofisher, Cat. #: 10569010), FBS (ThermoFisher, Cat. #: A3160701), Pen-Strep (10000 U/ml) (Thermofisher, 15140122), EGM-2 ™ Endothelial Growth Medium-2 BulletKit™ (Lonza, CC-3162), 0.05% Trypsin-EDTA (Cat. #: 25300062), Lonza’s MSCGMT human Mesenchymal Stem Cell Growth BulletKit (Catalog #: PT-3001), Matrigel (MG), Growth factor-free (Cat. #: 40230), Collagen, rat tail, (Cat. #: 354236), Luciferase assay system (Promega, Cat. #: E1500), 384-well round bottom, ultra-low attachment spheroid microplates (Corning, Cat. #:3830) was purchased from Corning. 2-well culture inserts were purchased from ibidi (Ibidi Cat. #: 80209). Aurum Total RNA Mini Kit (Cat. #: 7326820), DNase I (Cat. #: 7326828), iTaq Universal SYBR Green Supermix (Cat. #: 1725121), and iScript cDNA Synthesis Kit (Cat. #: 1708891) were purchased from Bio-Rad. Tissue Culture Treated 24-well plate (LPS Cat. #: 702001), 75-cm^2^ Polystyrene tissue Culture-Treated Flasks (LPS Cat. #:708003), 60 mm tissue culture treated dishes (LPS, Cat. #: 705001), 96-well Cell Culture Plate (LPS Cat. #: 701001), 6-well Cell Culture Plate (LPS Cat. #: 703001), 96-well PCR plates (LPS, Cat. #: L223080), PCR Plate Covers (LPS Cat. #: HOTS-100) were purchased from Laboratory Products Sales (LPS). Vybrant DiD Cell-Labeling Solution (Thermofisher, Cat. #: V22887), Vybrant DiO Cell-Labeling Solution (Thermofisher, Cat. #: V22886), Vybrant Dil Cell-Labeling Solution (Thermofisher, Cat. #: V22885) were purchased from Thermofisher. Thrombin, human plasma-derived (Sigma, Cat. No #: T6884-100UN) and Fibronogen, human plasma-derived (Sigma, Cat. No #: F4883-500MG) were purchased from Sigma Aldrich. All PCR primers were purchased from either Integrated DNA technologies (IDT), Sigma Aldrich, or Thermofisher.

### Cell lines

HepG2 liver carcinoma cells (ATCC®, Cat. #: HB-8065), MRC-5 lung fibroblasts cells (ATCC®, Cat. #: CCL-171), and Human embryonic kidney cells (HEK-293) (ATCC®, Cat. #: CRL-3216) were purchased from ATCC. Bone marrow mesenchymal stem cells (BM-MSC) (Lonza®, Cat. #: PT-2501), and Human umbilical vein endothelial cells (HUVEC) (Lonza®, Cat. #CC-2935) were purchased from Lonza (Basel, Switzerland). Human foreskin fibroblasts (HFF) cells was a kind gift from Professor Stelios Andreadis (University at Buffalo),

### Antibodies

Mouse anti-human albumin monoclonal antibody (sc-271604), Mouse anti-human FOXA2 monoclonal antibody (sc-374376) and mouse anti-human AFP monoclonal antibody (sc-130302) were purchased from Santa Cruz Biotechnology. Ki-67 recombinant rabbit monoclonal antibody (SP6) (Thermofisher, Cat. #: MA5-14520), Goat anti-Mouse IgG (H+L) Cross-Adsorbed Secondary Antibody, Alexa Fluor 488 (Thermofisher, Cat. #: A-11001), Goat anti-Rabbit IgG (H+L) Cross-Adsorbed Secondary Antibody, Alexa Fluor 594 (Thermofisher, Cat. #: A-11012) were purchased from Thermofisher (Waltham, MA).

### Engineering a stable GFP/firefly luciferase HepG2 cell line

HepG2 cells (p10) cultivated at 200,000 cells/well in a 6-well plate, were transfected with a third generation lentivirus bearing a transgene (ubiquitin-fluc2-egfp (Human Ubiquitin promoter, Firefly luciferase 2, enhanced green fluorescent protein)), as done previously, (Parashurama, Ahn et al. 2016) at an MOI of 5. To select for stably expressing cells, HepG2-Fluc2-eGFP cells were expanded, sorted by fluorescence activated cell sorting for high expressing cells, and replated, and then serial sorted 2 more times over a 4-week period, expanded, frozen in aliquots, and used in all experiments.

### Barrier migration assay

Briefly, HepG2 cells were seeded into a 2 chamber Ibidi co-culture system. This system consisted of 2 80 µm/cm^2^ regions separated by a 500 µm gap. 80,000 HepG2 cells were seeded into each region. After cells settled overnight, the barrier was removed and cells were allowed to migrate over time. Images were captured every 30 minutes over a 24-hour time period using phase contrast microscopy. Images were captured at 10x magnification.

### Transwell cell migration

Cell migration was determined using 8 µm pore size transwell (Corning Incorporated) made to fit into 24 well plates. Briefly 100,000 HepG2 cells expressing GFP/fluc, were seeded onto the upper membrane of the Transwell. MRC-5 fibroblasts were added at varying concentrations was added to the bottom well to serve as chemoattractant. Following incubation for 24 hours at 37°C, non-invading cells remaining on the upper surface were removed using a wet cotton swab, whereas cells on the lower surface were imaged using fluorescence microscopy for GFP expression of cells that had migrated. Migrating cells were subsequently analyzed and quantified. Images obtained at 10x magnification.

### Preparation of agarose-coated microwells

1 wt% agarose is made by dissolving 1 gram of agarose powder into 100 mL of room temperature distilled water. The mixture is then autoclaved to ensure sterility. To create round bottom wells the agarose hydrogel is warmed until in liquid phase and then 50 µL is pipetted into each individual well of a 96-well plate (Corning). The plate is allowed to cool at room temperature for 20 minutes before cells and media are seeded into it. These coated microwells are used therefore to form the epithelial spheroids. We observed no change in morphology or gene expression on agarose-coated plates.

### Spheroid formation assay

HepG2 and HEK 293-T cells were grown in a T-75 flask with DMEM supplemented by 10% FBS and 1% Pen-strep. Cells were maintained in an incubator operating at 37°C and 5% CO_2_, and the growth medium was changed every other day. The cells were cultured to 80-90% confluence and then passaged by aspirating the used growth medium from the T-75 flask and rinsing with 1x PBS. The flask was then filled with 5 mL of 0.05% Trypsin-EDTA solution and left to incubate for 5-10 minutes at 37°C. Next, the mixture of trypsin and the cells was then aspirated into a 15 mL centrifuge tube and filled with 5 mL of complete growth medium to inhibit trypsin activity. To wash the cells, the tube was centrifuged at 300 g for 5 minutes and the supernatant was discarded and replaced with PBS. The cell pellet at the bottom of the tube was re-suspended by pipetting. The tube was then centrifuged again at 300 g for 5 minutes and the supernatant was discarded. The cells were then re-suspended in complete growth medium (DMEM containing 10% FBS and 1% Pen-strep) and diluted to a final concentration of 10,000 cells per 1 mL of growth medium. Cells were then mixed each time, before pipetting 100 µL into a number of wells of a sterile Agarose coated 96-well plate. Parafilm was wrapped around the outside of the plate to secure the lid and to ensure sterility, and the plate was centrifuged at 1000 rpm for 5 minutes to bring the cells to the center, bottom of the wells. Plates were incubated in 37°C and 5% CO2 for 72 hours until spheroids formed and imaged after each 24 hours using phase contrast and fluorescent microscopy. Spheroids were further processed depending on the application.

### Fluorescent dye-labeling of cell lines

Briefly, cell lines are harvested using 0.05% trypsin and collected in suspension, counted and adjusted to a final concentration of 1×10^6^ cells/mL in a microcentrifuge tube. Then 5 uL of either Vybrant DiD (red), Dil (yellow), or DiO (green) cell-labeling solution respectively is added to each vial containing cells. The cell/dye mixture is then incubated for 20 minutes at 37°C before being centrifuged at 1500 rpm for 5 minutes to separate the supernatant containing residual dye. The remaining cell pellet is then washed twice in fresh DMEM before being seeded in ultra-low attachment round bottom plates to generate fluorescently labeled cells which can be used for cell culture or spheroid culture.

### Spheroid cultivation in extracellular matrices

HepG2 spheroids grown in 384- or 96-well, agarose-coated plates were allowed to form spheroids in suspension. Subsequently half the media inside each well was aspirated and an equal volume of matrix was introduced into each well for a 1:1 dilution. Additionally, pre-mixed MG/Collagen blends (mixed on ice) could also be introduced into each well. Collagen (2 mg/mL) was prepared as suggested by the manufacturer. Briefly, to make Collagen gels, rat tail Collagen was diluted with pre-calculated amounts of de-ionized water, 10x PBS, and 1N NaOH on ice, for a final concentration of 2 mg/ml. Fibrin hydrogels were made by pre-mixing fibrinogen (3.25 mg/mL), and thrombin (12.5 Units/mL) in a 4:1 ration on ice. The final solution contains equal amount of fibrinogen and thrombin, thus forming fibrin. The gels polymerize quickly at room temperature for spheroid embedding applications. Care was taken to pipette slowly into each well, to prevent injury to HepG2 spheroids. For experimental models with cells within the mesenchyme, BM-MSCs, HFF, MRC-5 cells, HUVECS, HEK cells were pre-mixed with a matrix component before being added as 20,000 cells/well to each well. Matrix embedding of HepG2 spheroids was performed on ice to minimize polymerization during seeding. Subsequently, plates were incubated for 30 minutes to ensure proper gel solidification before additional fresh serum containing DMEM was added to each well. Migration was then observed over a 10-day period.

### Spheroid MG outgrowth assay

A MG coated 6-well plate was made by seeding 800 uL/well of diluted (1:14 diluted) MG onto 6-well plates. To solidify the gel the plate was then allowed to incubate for 30 minutes. Separately, HepG2 spheroids were collected from 96-well plate culture and placed into a 15 mL conical tube. The spheroids were then rinsed once in fresh serum containing DMEM before being seeded as 15 spheroids per well of the matrix coated 6-well plates. Spheroids were allowed to attach to the plate overnight before cell conditioned media was introduced separately into each well. Spheroid migration was then observed over a 24-hour period.

### Mixed spheroid formation

HepG2 and HFF cells are mixed together in a 1:1 ratio (20,000 cells total) to form compact organoids in agarose coated 96-well plates. Spheroids are formed over a 24-hour period and are stable in culture. These multicellular spheroids are then embedded in different matrices to observe migration over time. To determine uniform presence of different cells, HepG2 and HFF cells are also pre-dye labeled (Vybrant DiD cell labeling solution) according to the manufacturer and then imaged.

### Preparation of conditioned medium from cell lines

Cell lines are seeded into a T-175 tissue culture treated (TCT) flask at a seeding density of 15,000 cells/cm^2^. Cells are allowed to adhere to the plate before media is replaced with 10 mL of culture media DMEM (FBS/Pens-strep). Cells are incubated for 48 hours before the media is collected in a 15 mL tube. The media is centrifuged at 1200 rpm for 5 minutes to allow for removal of cellular debris/dead cells. The media was then filtered through a 0.2 µm filter to ensure sterility. The media is then added to the matrix encapsulated HepG2 spheroids.

### Spheroid MG droplet migration assay

HepG2 and HEK spheroids were harvested from the 96-well plate and collected into a 15 mL conical tube. The spheroids were allowed to settle to the bottom of the dish based on gravity. After 10 minutes, spheroids were washed once with PBS before being resuspended in MG (GF-) diluted by half. 15 µL was then used to collect one spheroid at a time with MG and seeded onto a tissue culture treated (TCT) 60 mm dish. MG droplets were allowed to solidify in the incubator for 30 minutes. Afterwards conditioned media from MRC-5 fibroblasts was added to the dish. Spheroids were allowed to incubate for 4 days before significant migratory phenotype was observed.

### Spheroid Collagen gel droplet migration assay

The collagen droplet assay was performed similar to the MG droplet assay. Rat tail collagen type 1 matrix was diluted to 2 mg/mL according to the manufacturer. HepG2 spheroids were collected from 3D suspension in agarose coated plates and mixed with collagen type 1 matrix (2 mg/mL) in a 15 mL tube on ice. Then 15 µL droplets containing individual spheroids were seeded onto 60 mm tissue culture treated dishes. The droplets were allowed to gel in an incubator (20% O_2_, 37 °C) for 30 minutes prior to introducing conditioned media to the dishes. Spheroid migration was observed over a 4-day period.

### Diluted conditioned medium assay

MRC-5 conditioned media (CM) purified as explained previously was diluted by various ratios using fresh complete DMEM (10% growth serum, 1% pen-strep) in separate 15 mL tubes. Diluted conditioned medias of 1 CM:1 DMEM and 1 CM:7 DMEM were prepared and introduced to the matrix embedded spheroids. The migration of the spheroids was then observed over time (4 days).

### MRC-5 spheroid formation assay

MRC-5 cells were grown in a T-75 flask with DMEM supplemented by 10% FBS and 1% Pen-strep. Cells were maintained in an incubator operating at 37°C and 5% CO_2_, and the growth medium was changed as necessary. The cells were cultured to 80-90% confluence and then passaged by aspirating the used growth medium from the T-75 flask and rinsing with 1x PBS. The flask was then filled with 5 mL of 0.05% Trypsin-EDTA solution and left to incubate for 5-10 minutes at 37°C. The Trypsin-EDTA and the cells were then aspirated into a 10 mL centrifuge tube and filled with 5 mL of complete growth medium to stop the trypsinization. The tube was centrifuged at 300 g for 5 minutes and the supernatant was discarded and replaced with PBS. The cell pellet at the bottom of the tube was re-suspended by pipetting rigorously up and down. The tube was then centrifuged again at 300 g for 5 minutes and the supernatant was discarded. The cells were then re-suspended in complete growth medium and diluted to a final concentration of 2 x 10^5^ cells per 1 mL of growth medium. Cells were then mixed each time before seeding 50 µL per well of a 384-well ultra-low attachment plate. Parafilm was wrapped around the outside of the plate to secure the lid and to ensure sterility, and the plate was centrifuged at 1000 rpm for 5 minutes to bring the cells to the center, bottom of the wells. Plates were incubated in 37°C, and 5% CO_2_ for 24 hours until spheroids formed and then imaged using phase contrast and fluorescent microscopy. Spheroids were further processed depending on the application.

### Spheroids coculture in MG assay (384-well plate)

HepG2 spheroids were harvested into a 15 mL conical tube. The spheroids were incubated for 5-10 minutes to settle to the bottom of the tube before the medium was aspirated and disposed of. The spheroids were resuspended in fresh DMEM. 10 µL droplets containing a spheroid were then added to the pre-existing MRC-5 spheroids in a 384-well plate, one spheroid at a time; ensuring that each well contains both kinds of spheroids. 40 µL of MG (GF-) were then added to each well, resulting in a 1:5 dilution. The plate was incubated for 15 minutes to allow gel formation. 50 µL of fresh DMEM was then added to each well. Spheroid migration was observed over a 4-day period.

### Microscopy

For cellular imaging spheroids in various formats were imaged during morphogenesis using both phase contrast and fluorescence microscopy. Images were obtained using a Zeiss Axio fluorescence microscope.

### Histology

Spheroids were harvested from the ULA plates and fixed in 4% paraformaldehyde for 30 minutes at room temperature. They were then collected and embedded in agarose (2 wt %) prior to paraffin embedding. Paraffin embedded blocks were then sectioned at 10 µm. Antigen retrieval was performed by heating rehydrated section in 1x Tris-EDTA buffer solutions for 20 mins in microwave. Slides were then used for subsequent staining. Paraffin embedded 10 µm sections were stained with Eosin and Hematoxylin (Eosin Y Catalog Number (DcE-40), Hematoxylin Catalog Number (DcH-48)) and mounted with medium before microscopy.

### Immunostaining of monolayer and 3D spheroid culture

For detection of intracellular localization of Albumin, AFP, FOXA2, or Ki-67, cells cultured on tissue culture treated dishes were washed once with PBS and then fixed with 4% paraformaldehyde, permeabilized with 0.1% Triton X-100 in PBS and blocked with 1% BSA in PBS for 30 minutes. Following this, plates were incubated with primary antibodies overnight and then rinsed before incubation with secondary antibody (AlexaFlour 488, Thermofisher) for one hour. Subsequently plates were incubated with DAPI (4,6-diamidino-2-phenylindole) for nuclear detection for 10 mins at room temperature. An additional staining protocol was developed and optimized to stain whole mount HepG2 spheroids in suspension. Briefly, HepG2 organoids were fixed in 4% PFA for 1 hour and then blocked for 2 hours in 1% BSA. The spheroids were then incubated with primary antibody at 5 ug/mL with gentle agitation at 4°C overnight. The following day, the organoids are washed 3 times with 1% PBST (each wash 20 mins) under gentle agitation at room temperature. Secondary antibody (AlexaFlour 488, Thermofisher,) was then added for incubation at 4°C overnight and washed out as described above. DAPI incubation (10 minutes) was used to counterstain before images were obtained. For immunolabeling of spheroids that generated protrusions in the presence of MRC-5 CM, the same protocol for organoid staining was used. Ki-67 monoclonal primary antibody was used to stain the migrating spheroids overnight at 4°C. The following day, secondary antibody (AlexaFlour 594 Thermofisher) was applied again for an additional incubation overnight at 4°C before DAPI incubation (10 minutes) and subsequent imaging.

### Fluorescence microscopy

Cells were visualized under fluorescence microscopy using a standard filter for green, blue or red fluorescence. Images were captured using a Zeiss Axio fluorescence microscope.

### Reverse transcriptase polymerase chain reaction (RT-PCR)

For each experimental condition, cell lysates were collected using the Aurum Total RNA Mini Kit. Total RNA was isolated from duplicate or triplicate samples and concentrations were measured using the NanoDrop One/One Microvolume UV-Vis Spectrophotometer (Thermofisher). CDNA was synthesized using the iScript cDNA synthesis kit (Biorad) and made using an Eppendorf 5331 MasterCycler Gradient Thermal Cycler (Eppendorf) with 5ng of RNA for each planned qRT-PCR reaction. Each sample was plated in triplicate in a reaction volume consisting of 10 µL per well. Each well had 5 µL of iTaq™ Universal SYBR® Green Supermix, and forward and reverse primers at a concentration of 300nM. The qRT-PCR reactions were run for 40 cycles in a Biorad C1000 Touch Thermal Cycler. Gene expression analysis was conducted utilizing the delta-delta-Ct method, with GAPDH/B-actin used as normalization housekeeping gene(s). Primer sequences are reported in Supplementary Information.

### Analysis of spheroid migration

ImageJ was used to determine relative properties of the migrating spheroids. Images were uploaded to ImageJ, with a pre-set scale used to quantify images taken at the same objective focus. The length feature in Image J was used to determine protrusion length and thickness in the various experimental conditions. The count plugin feature was used to determine the number of cords or emerging strands in different fields of view and over time. The trace plugin feature in ImageJ was used to outline the edge of the spheroids over time to estimate the growth of the spheroid. The Skeletonize3D plugin in image J was used to analyze the branching phenotype observed in migrating spheroids. Specifically for analysis in Skeletonize3D plugin, images were first converted to 8 bit grey-scale, a threshold was applied to isolate only the branching at the edge of the spheroid before Skeletonize 3D was used to carry out branching analysis.

### Statistics

Data collected was reviewed and analyzed by GraphPad Prism (version 7) or Microsoft excel. Student’s t test was used to determine statistical differences between two independent groups (P value set at <0.05).

## RESULTS

### Mesenchymal stem cells induce liver cell migration, but not in 3D models of the liver diverticulum

We began modeling the liver diverticulum (Fig. 1 A-B) by employing a well-established, stable liver cell line (HepG2) which expressed an immature hepatocyte gene signature, and human mesenchymal stem cells (hMSC). hMSC and human umbilical vein endothelial cells (HUVEC) have been used to model the liver bud (E10.0) but not the liver diverticulum (E8.5) (Takebe, Sekine et al. 2017). Based on the hypothesis that mesenchyme can induce factors that elicit liver cell migration, we assessed migration potential employing a commercially available wound healing model (Ibidi 2-well culture insert). The system contained two islands for cell adhesion separated by a 500 µm gap. In the presence of stable liver cells on each island, there was no changes observed over 24 hours. However, when hMSC or HUVEC were added to the second island, we observed significant changes in both border closure rate, for both HUVEC and MSC, and liver-specific border area in the MSC condition only, over 24 hours (Fig. 1C-D). The data suggests that MSC can induce liver cell migration. Next, we performed a trans-well assay, and liver cells migrated towards the MSC, demonstrating a statistically significant, induction of migration compared to controls (Fig. 1E-F). We wanted to determine if this affect was present in a 3D model since mechanisms of 2D and 3D differ. We engineered a model of the liver diverticulum by generating liver spheroids of a specific size (Fig. 1G-H). We demonstrated that spheroids form uniformly (Fig. 1H) and cell dilution/ labeling studies demonstrated that multiple clones give rise to the spheroid without aggregation, suggesting that clones grow equally and aggregation of separate clusters and clonal dominance are not the main mechanisms of spheroid formation (Fig. 1I). We performed gene expression analysis for liver differentiation genes to compare conventional and 3D culture. We found a decrease in Foxa2 expression, and otherwise no significant changes (Fig. 1J). We performed whole spheroid immunostaining to demonstrate homogenous albumin (Alb), alpha-fetoprotein (AFP), and Foxa2 expression (Fig. 1K, Supp. Fig 1), and tissue histology demonstrating no central necrosis by day 3 (Fig. 1L). Using these 3D spheroids, we modeled the liver diverticulum by modeling the STM with MG seeded with 20,000 hMSC. Liver cells were engineered with eGFP, spheroids were formed for 3 days, and hMSC were labeled with cell membrane dye Vybrant DiD (red) (Fig. 1M). Interestingly, within 24 hours and over a 3-day period, hMSC migrated towards the liver spheroids (Fig. 1N). Further, we observed no liver spheroid migration into the mesenchyme, and alternatively, significant migration of MSC towards liver spheroids (Fig. 1O). We tested several conditions of gel formation and observed the same effect (data not shown), indicating this was not due gelation conditions. We also tested liver/HUVEC based on previous literature (Pettinato, Lehoux et al. 2019); (Jung, Kang et al. 2017) but we observed that HUVEC themselves aggregated within MG and then aggregated with the edge of the sphere, with no uniform liver cell migration (Supp. Fig. 2). We conclude that both MSC or HUVEC may not be the ideal within MG to induce liver cell migration in this model of the liver diverticulum.

**Figure 1.**
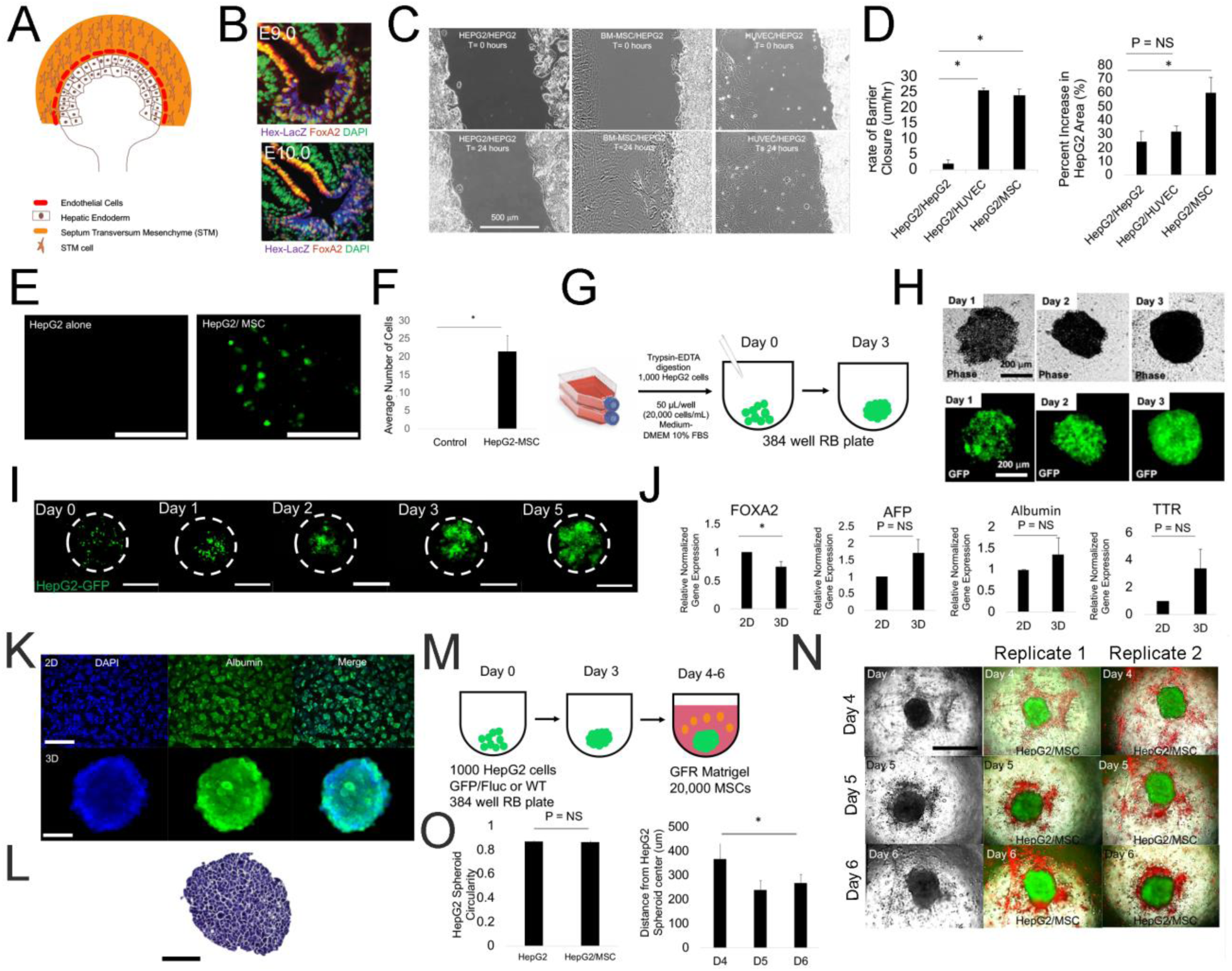
Mesenchymal stem cells induce liver cell migration, but not in 3D models of the liver diverticulum. A) Schematic of liver diverticulum (LD) during mouse liver development (E8.5). Shown are epithelial hepatic progenitor cells (brown), endothelial precursors (red) and the septum transversum mesenchyme (STM) (orange). At E9.5, hepatic progenitor delaminates from basement membrane, and together with endothelial cells, co-migrates and interacts with STM and STM cells, forming the hepatic cords. B) Hepatic cords identified in the embryological liver (E9.0, top) give rise to hepatic cords migrating (E9.5, bottom) with evidence of co-migration, branching, and interstitial migration, marked by Hex and Foxa2. Figure adopted with permission from (Bort, Signore et al. 2006). C) Phase contrast images of 2D barrier migration assays between HepG2 and mesenchymal stem cells (HUVECS and MSCs) on at T=0 hours (top) and T = 24 hours (bottom). Time lapse performed every 30 minutes over 24 hours. D) Left- Bar graph analysis of migration demonstrates a significant increase in barrier closure in the presence of HUVECS (P = 0.00057, n = 3) and MSCs (P = 0.00028, n = 3) as compared to HepG2 control (n = 3). Right- Bar graph analysis of HepG2 border wall movement in the presence of MSCs (P= 0.047, n = 3) as compared to HepG2 control (n = 3). Plotted is mean ± SD. Significance defined as P ≤ 0.05. E) Fluorescence image focused on a transwell during a transwell assay demonstrating HepG2-GFP migration alone (left) or in the presence of MSCs (right). Bar = 200 µm. F) Bar graph plotting average number of HepG2-GFP migrating through transwell (P = 0.013, n=3). Plotted is mean ± SD. Significance defined as P ≤ 0.05. G) Schematic for making spheroid from HepG2 or HepG2-GFP cells. 1000 cells are added in 50 µl and to a 384-well round bottom non-adherent plate, and cultured for 3 days. H) Phase contrast (top) and corresponding fluorescent images (below) of HepG2-GFP/HepG2 (1:10) mixed spheroids on Days 0-5 during spheroid formation. I) Fluorescent images of dilution experiments of HepG2-GFP/HepG2 (1:10) mixed spheroids on Days 0-5 during spheroid formation. Bar = 300 µm. J) qRT-PCR analysis of Foxa2 (P = 0.040940714, n = 3), AFP, albumin, and TTR (P = NS) in monolayer and day 3 spheroid culture compared to control (n=3). Plotted is mean ± SD. Significance defined as P ≤ 0.05. K) Fluorescent images of DAPI (nucleus) and albumin in Hep2 monolayer (top) and spheroid (bottom) culture on day 3 of culture. L) Hematoxylin and eosin stained 10 µm section of HepG2 spheroid in suspension culture illustrates uniform epithelial morphology of 3D tissue. Bar = 200 µm. M) Schematic for modeling the liver diverticulum. Spheroids are made as in (G) and then embedded in 20,000 MSC containing growth factor-free (GFR) Matrigel (MG) and cultured for long term. N) Phase (left), double fluorescent (red/green) images (right) of day 4, 5, and 6 liver diverticulum models, bearing a HepG2-GFP spheroid (green) and MSC (red) in MG. Right- Columns of images are Replicates 1 and 2 with double fluorescent images on Days 4, 5, 6. Bar = 500 µm. O) Bar graph plotting spheroid circularity (left) of HepG2 and HepG2/MSC spheroids (P = NS, n = 3) and (MSC) distance from HepG2 spheroid on days 4, 5, and 6. Day 4 to 6 compared (P = 0.00043, n=3). Plotted is mean ± SD. Significance defined as P ≤ 0.05.

**Figure 2.**
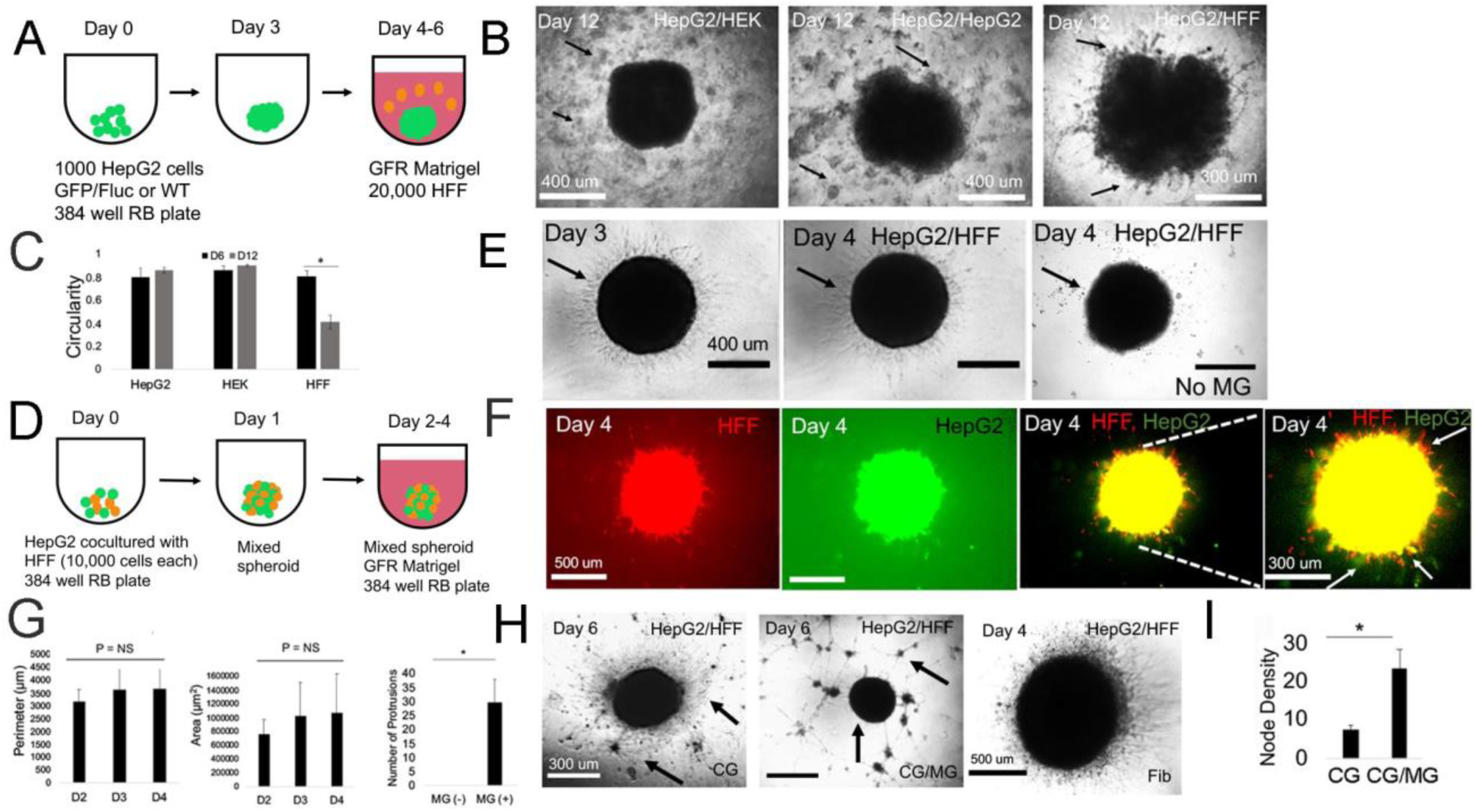
Mixed spheroids with fibroblasts result in 3D liver collective cell migration. A) Schematic for modeling the liver diverticulum. HepG2 spheroids are embedded in 20,000 human foreskin fibroblasts (HFF) containing growth factor-free (GFR) Matrigel (MG) and cultured for long term. B) Phase contrast images on Day 12 of culture of negative control (HepG2 spheroid embedded with human embryonic kidney (HEK) cells (HepG2/HEK)), a negative control (HepG2/HepG2), and the experimental (HepG2/HFF) condition all at 1:1 ratio. Arrows in the HepG2/HepG2 condition and the HepG2/HEK condition specify the relative location of the HepG2 and HEK cells seeded in the surrounding GFR matrigel. Arrows in the HFF condition specify the occurrence of co-migration of both HepG2 and HFF cells into the MG. C) Bar graph plotting spheroid circularity in HepG2/HepG2, HepG2/HEK, HepG2/HFF spheroids. Comparison of HepG2/HFF shown (P = 0.0000343, n=3). Plotted is mean ± SD. Significance defined as P ≤ 0.05. D) Schematic for modeling the liver diverticulum with mixed spheroid containing HepG2 cells and HFF cells cultivated for 1 day followed by embedding in MG from Days 2-4. E) Phase contrast images of HepG2-GFP/HFF mixed spheroids in MG on Day 3 and 4, and Day 4 no MG (negative control) condition. Arrows in the HepG2/HFF condition migration of HFF cells into the MG. In the control condition containing no MG, arrows denote lack of migration. F) Fluorescent images of Day 4 HepG2-GFP/HFF mixed spheroids in MG. From Left to right- HFF (red) cells. HepG2-GFP (green), HFF/HepG2 (red, green), HFF/HepG2 (high magnification). White arrows show red, yellow, and green projections. G) Bar graph plotting perimeter (left), area (middle) of Day 4 HepG2-GFP/HFF mixed spheroids over time. P = NS (n = 3). Right- Bar graph plotting number of protrusions in the HepG2/HFF spheroids on Day 4 in without MG (MG -) and with MG (MG +), (P = 0.022, n=3). Plotted is mean ± SD. Significance defined as P ≤ 0.05. H) Phase contrast images of HepG2/HFF mixed spheroids. Left- in collagen gel (CG) on day 6 arrows who hair-like structures migrating. Middle- in CG/ MG mixture, arrows show node like structures in hydrogel. Right- in Fibrin hydrogel, hair-like structures are migrating. I) Bar graph plotting Node density within the mesenchyme, comparing HepG2/HFF spheroids grown in the CG and CG/MG condition (Day 6, P = 0.00427, n = 3). Significance defined as P ≤ 0.05.

### Mixed spheroids with fibroblasts result in 3D liver collective cell migration

We wanted to induce liver spheroid migration in our liver diverticulum model. We found that human foreskin fibroblasts (HFF) not only migrated towards liver cells, but they also induced liver cell migration in 2D models (Supp. Fig. 3). We employed MG with human foreskin fibroblasts (HFF), as an alternative to MSC or HUVEC (Fig. 2A). Compared to controls, we observed evidence of perpendicular, cellular protrusions and cellular strands at the edges of the liver spheroid in the presence of the HFF, which was statistically much higher than controls, by day 12 (Fig. 2B-C). However, it was unclear whether HFF were joining the spheroid and were part of the migrating strands, and it took a long time to observe the effect. We re-examined the liver diverticulum, and when it forms the liver bud, intermingling between hepatic endoderm and STM cells occurs (Fig. 1). We hypothesized that creating mixed spheroid coculture, that mimic the intermingling in the liver bud, may enhance the kinetics of 3D CCF formation. Compared to controls, we observed rapid compaction of the spheroids (24 hours vs 3 days) (Fig. 2D). To determine individual cell fate within the mixed spheroid, we performed double labeling of liver (eGFP) and HFF (DiI dye) cells after embedding in MG. Serial *in vitro* microscopy demonstrated uniform, perpendicular strands that increase with time (Fig. 2E). To determine if co-migration occurred, in which both HFF and liver cells migrate together, we imaged the cells after fluorescent labeling, performed single well fluorescent imaging and adjusted the images at high intensity to clearly visualize migrating strands. The data demonstrated a large number of migrating strands, including red strands, yellow strands (both green and red), and in a few cases, green strands (Fig. 2F). Our analysis indicated statistical differences between presence and absence of MG (Fig. 2G). Next, we tested the effects of the extracellular matrix composition. We found that with culture of mixed spheroids of collagen gels, fibroblasts appear to migrate in an elongated fashion extensively into the matrix (Fig. 2H, **left**). In fibrin gels, both fibroblasts and HepG2 cells individually take an elongated morphology (Fig. 2H, **middle**, Supp. Fig. 3), while in collagen/MG mixtures, we observe fibroblast migration and statistically significant nodule formation (Fig. 2H, **right, 2I**). Despite the varying morphogenesis we observed when we varied matrix composition, we don’t observe 3D CCM of liver cells in the collagen, collagen/MG, and fibrin conditions, suggesting that they did not play a major role in co-migration during liver CCM from mixed spheroids.

**Figure 3.**
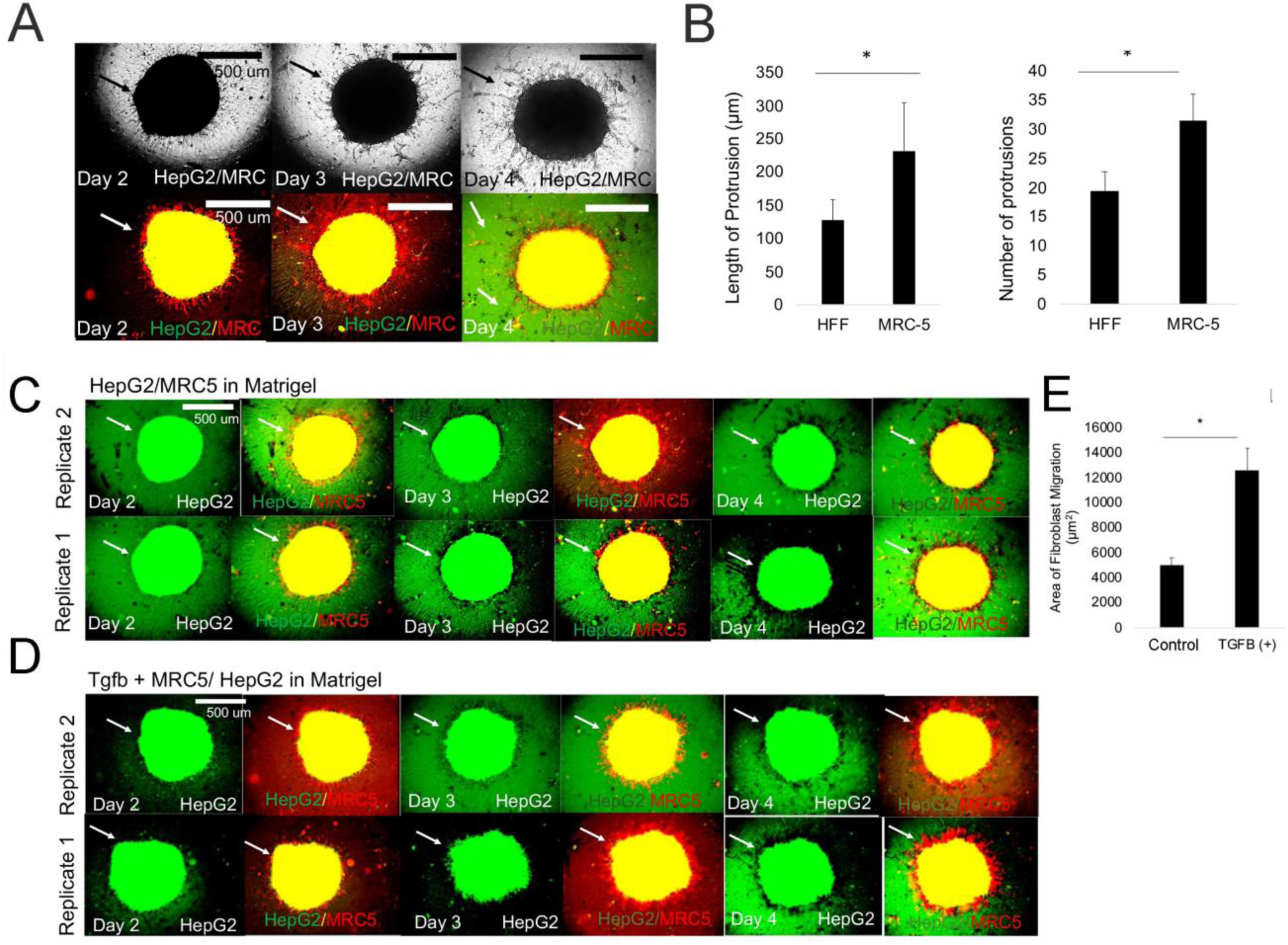
MRC-induced fibroblasts enhance 3D liver collective cell migration in a mixed spheroid model. A) Phase contrast (above) and fluorescent images (red, green) below of HepG2-GFP/MRC-5 mixed spheroids in MG on day 2, 3 and 4, and a day 4. Arrow in the picture specifies emergence of finger like protrusions. B) Bar graph comparing length of protrusions (left) (P = 0.0076, n = 3), number of protrusions (right) (P = 0.033, right, n=3) on day 2, 3, and 4. Plotted is mean ± SD. Significance defined as P ≤ 0.05. C) Fluorescent images of Day 2-4 of HepG2-GFP/MRC-5 mixed spheroids in MG. From Left to right- HepG2 (green) cells, combined HepG2 (red) and MRC-5 (yellow) images. Replicates 1 (above) and 2 (below) are shown. Arrows show HepG2 and MRC-5 migration. D) Fluorescent images of Day 2-4 of HepG2-GFP/MRC-5 mixed spheroids in MG after treatment with TGFß1 (20 ng/mL). From Left to Right- HepG2 (green) cells, combined HepG2 (red) and MRC-5 (yellow) images. Replicates 1 (above) and 2 (below) are shown. Arrows show HepG2 and MRC-5 migration. E) Bar graph comparing area of fibroblasts migration in negative control (HepG2-GFP/MRC-5) and (HepG2-GFP/MRC-5 + TGFß1 20 ng/mL), P = 0.011617, n= 3. Plotted is mean ± SD. Significance defined as P ≤ 0.05.

### MRC-induced fibroblasts enhance 3D liver collective cell migration in a mixed spheroid model

MRC-5 fibroblasts are derived from human fetal lung tissue and unique amongst human fibroblast lines to induce epithelial cell scattering reduce junctional connections in 2D *in vitro* models (Stoker and Perryman 1985). Given that the STM is composed of fetal mesenchyme, and that the lung and the liver are from a common precursor, we hypothesized that MRC-5 fibroblasts may be a useful tool to model the STM cells and induce liver 3D CCM. We observed significantly more migration (length and number of protrusions) in the case of MRC-5 condition on day 3 compared to the HFF condition (**compare** Fig. 2E **to** Fig 3A). Since TGFß pathway potently effects hepatic cord formation, we wanted to evaluate if 3D CCM is affected by this pathway. When we added TGFß1 growth factor to mixed spheroids with MRC-5 cells, we observed an increase in the projected area of migration in the MRC-5 condition compared to the control condition (**compare** Fig. 3B to 3D). These experiments suggest that MRC-5 cells specifically enhance liver 3D CCM.

### MRC-5-conditioned medium induces 3D collective cell migration within the *in vitro* liver diverticulum model

While liver and MRC-5 mixed spheroids did result in evidence of co-migration (both red and green), we wanted to further understand how MRC-5 resulted in an enhanced 3D liver CCM. We evaluated MRC-5- conditioned medium, as MRC-5 conditioned medium has been previously been shown to increase liver cancer cell migration in trans-well assays (Ding, Chen et al. 2015), but not for 3D migration. We first performed *in vitro* migration assays employing the Ibidi system as above (Fig. 1). The addition of MRC-5- conditioned medium resulted in statistically significant HepG2 border migration compared to conventional medium change (Fig. 4A-B). Next, we performed a trans-well assay, in which we tested liver cell migration in response to conditioned medium from MRC-5 fibroblasts (M-CM). We found a statistically significant increase in migration of liver cells in response to MRC-5 cells compared to hMSC (Fig. 4C). TO determine whether this effect was specific to MRC5, we performed a classic outgrowth assay in which liver spheroids were cultured in control medium, HepG2-conditioned medium, HFF-conditioned medium, and M-CM. The data demonstrated clear, statistically significant increase in spheroid outgrowth in the presence of MRC-5- conditioned conditioned medium after 24 hours of outgrowth (Fig. 4E-F). The data also demonstrated a down regulation of E-Cadherin at the edge of the spheroid (Supp. Fig. 5). This outgrowth assay demonstrated migration on a tissue plastic, but strictly this did not represent 3D migration. To test 3D migration, we developed a 3D migration assay by employing liver spheroids in MG droplets (Lancaster, Renner et al. 2013) in which spheroids remain suspended (Fig. 4G, Supp. Fig. 6). In this system, we first determined how the strength of MRC-5-conditioned medium affects 3D cell migration. We found that from day 4-11, MRC-5-conditioned medium M-CM led to 3D CCM (Compare Fig. 4H **(M-CM condition) to 4F (M-CM condition)**) clearly distinguishing it from a 2D outgrowth assay from the liver spheroid. To assess the mechanism by which M-CM functions, we performed dilution of conditioned medium (1:1, 1:7, M-CM first day only) and demonstrated statistically significant differences in 3D CCM as a function of dilution, as measured by area of growth, perimeter, protrusion, and cord count (Fig. 4H-K). When we diluted M-CM at a ratio of 1:19, we observed no evidence of migration (Supp. Fig. 7). Of note branching increases over time from day 4 to day 7 (Fig. 4J). We analyzed this 3D migration at high magnification to better understand morphogenetic processes, and demonstrate various aspects of migration. On day 7 in MG droplets, we observe sub-micron cellular strands including long strands with small branching, long strands without branching, thicker strands, and strands with multiple levels of branching (Fig. 4L**, Day 7**). Cell nuclei were not visible. However, when we performed nuclear (DAPI) and Ki67 staining on day 7 of spheroid culture (Supp. Fig. 8), we found nuclear staining within 3D protrusions. Further, we observed heterogeneous areas of Ki67 staining. On the other hand, by day 11 of culture, the migrating strands appeared to be interconnected, formed a sheet of cells, and attached to the dish, and the cells had spread with visible nuclei (Fig. 4L**, Day 11**). Even though the cells appeared to touch the dish, the cells exhibited branching and elongated morphologies on day 11, when compared to traditional outgrowth assay (**compare 4F to 4L**). We tested conditioned medium from other cell types and observed no effects, but observed that MRC-5 also induces 3D CCM in HEK cells by day 11 (Fig. 4M). To evaluate the effect of matrix on 3D migration, we tested the same liver spheroid model in collagen (CG) gels. Interestingly, by day 7 of culture, we observed highly linear, micron or submicron thick, cellular strands extending from the liver spheroid. We clearly observed linear strands with less branching than observed than in MG **compare 4L (Day 7) to 4N-O (Day 7)).** In collagen, we demonstrated a statistically significant increase in protrusion length and cord count with M-CM compared to control medium (Fig. 4P). When we compared the MG and CG conditions, we found that protrusion thickness was significantly reduced in the CG condition, and the number protrusions was comparable in both the CG and MG conditions. When we compared extent of branching on day 7, we found that CG had statistically less branching then MG condition (Fig 4P). We then developed a hypothesis that M-CM may induce migration via TGFß. We tested the TGFß pathway inhibitor A83-01, which inhibits TGFß, Activin and Nodal signaling pathway, by inhibiting ALK4, ALK5, ALK7 receptors and prevents SMAD2/3 phosphorylation. We observed a statistically significant and dose-sensitive inhibition of migration (Fig. 4Q-R). Collectively, these data demonstrate robust collective cell migration within the liver diverticulum model of MRC-5-conditioned medium.

**Figure 4.**
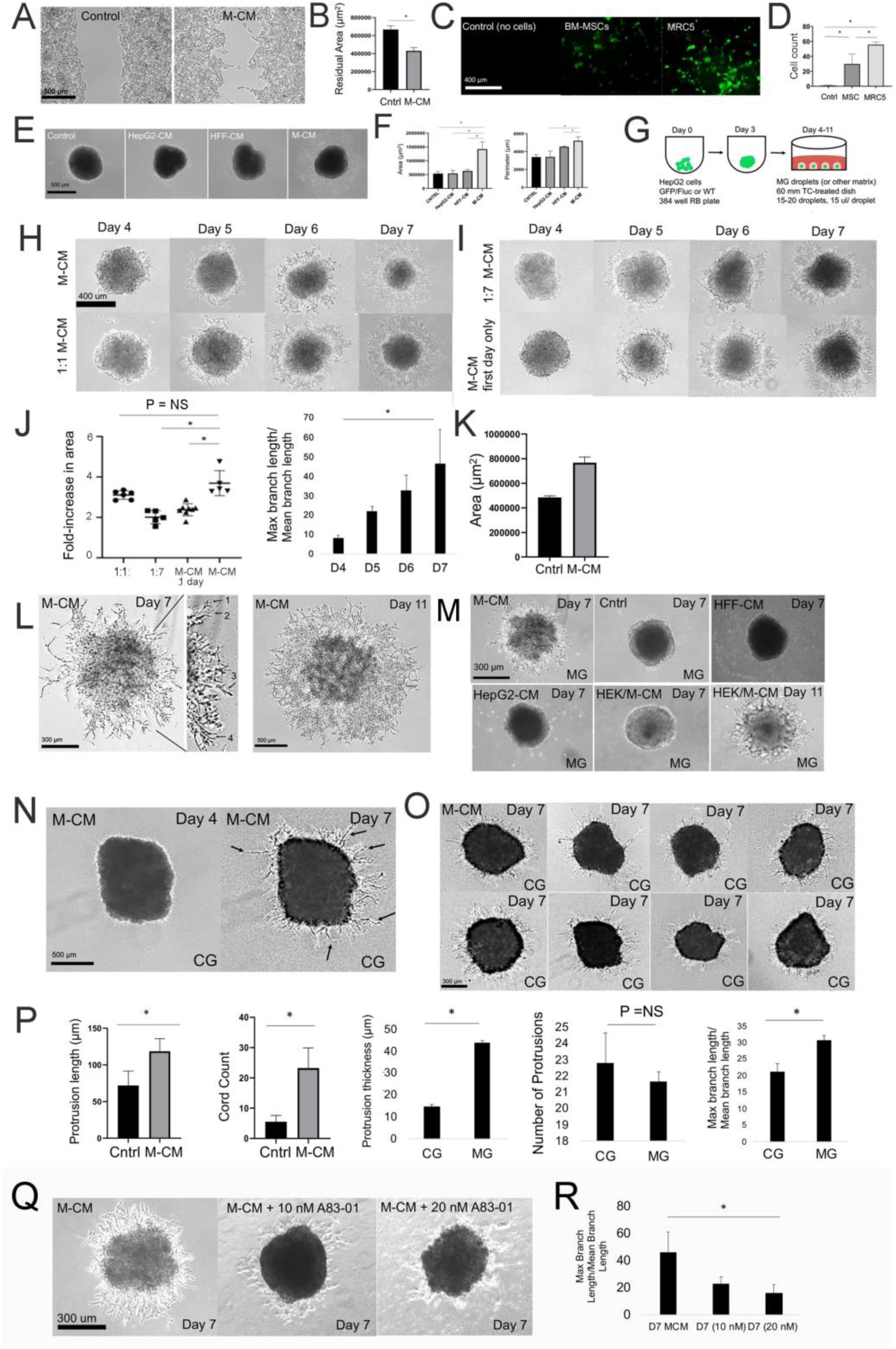
MRC-5-conditioned medium induces 3D collective cell migration within the *in vitro* liver diverticulum model. A) Phase contrast images of 2D barrier migration assays between HepG2 cells (on both sides of the barrier) in the presence of MRC-5-conditioned medium (M-CM) at T= 24 hours in control (left) and M-CM (right) conditions. B) Bar graph comparing residual area between control and M-CM conditions (P = 0.00085, n=3). Plotted is mean ± SD. Significance defined as P ≤ 0.05. C) Fluorescence image focused on a transwell during a transwell assay demonstrating HepG2-GFP migration alone, or in the presence of MRC-5 (and resulting M-CM), after 24 hours. D) Bar graph analyzing transwell assay comparing control to MSC (P= 0.023, n=3), MSC to MRC-5 MSC (P = 0.012, n = 3), and control to M-CM conditions (P = 0.0014, n=3). Plotted is mean ± SD. Significance defined as P ≤ 0.05. E) Phase contrast images of HepG2 spheroid outgrowth assay in control, HepG2 CM, HFF-CM, and M-CM (MRC-5 CM). Spheroids cultured on MG (1:15) coated plates. F) Bar graph for spheroid outgrowth assay comparing area of spheroid growth (left) and growth perimeter (right) between control, HepG2-CM, HFF-CM, M-CM, conditions. Comparison of spheroid growth area for M-CM and control (P = 0.0023, n=3), M-CM and HepG2-CM (P = 0.0030, n = 3), and M-CM and HFF-CM (P = 0.0029, n = 3) as shown. Comparison of growth perimeter between M-CM and HepG2-CM (P =0.0030, n = 3) and M-CM and HFF-CM (P =0.0029, n = 3) as shown. Plotted is mean ± SD. Significance defined as P ≤ 0.05. G) Schematic for modeling the liver diverticulum in MG droplets. HepG2 spheroids are embedded in MG droplets (15 µl) with 15-20 droplets per dish and cultured in M-CM medium. H-I) Phase contrast images HepG2 spheroids in MG droplet assay in study of M-CM dilution on Days 4 - Day 7. Conditions tested were Left-M-CM, 1:1 M-CM and Right-1:7 M-CM, M-CM first day only. For M-CM first day only, M-CM was added from Day 3-Day 4, washed gently, and then switched to M-CM. J) Analysis of MG droplet assay with M-CM. Left- Plot of fold-change in area across different M-CM dilutions (1:1, 1:7, M-CM one day only, and M-CM). Comparison of M-CM and 1:1 condition (P = 0.15, n = 3 for both conditions), M-CM and 1:7 condition (P = 0.0056, n = 3 for both conditions), and M-CM and M-CM one day only condition (P = 0.019, n = 3 for both conditions). Right- Measure of branching of morphogenesis (Max branch length/ Mean branch length) between Day 4- Day 7, comparison between Day 7 and Day 4 shown (P= 0.047, n=3). Plotted is mean ± SD. Significance defined as P ≤ 0.05. K) Bar graph comparing growth area of HepG2 spheroid in MG droplet assay in Control (no CM) and M-CM conditions on Day 7 of culture (P =0.010, n = 3). Plotted is mean ± SD. Significance defined as P ≤ 0.05. L) Phase contrast images of Day 7 (low and high magnification) and Day 11 liver spheroid in the MG droplet system cultivated with M-CM. Left- High magnification view demonstrates various stages/modes of 3D collective migration including 1- filopodia branching, 2- filopodia elongation 2- cell and nuclear extension (nucleus is visualized) 4- branches that are interacting. M) Phase contrast images on Day 7 of HepG2 spheroid in MG culture treated with various conditioned medium conditions including M-CM, HFF-CM, HepG2-CM, HEK-CM (Day 7 and 11 shown). N) Phase contrast images on Day 4 and Day 7 of HepG2 spheroid in Collagen (CG) droplet and M-CM medium. Arrows specify thin filopodia like extensions into collagen. O) Same as N except several experimental replicates shown on Day 7. P) Bar graph analysis of HepG2 spheroids in CG in control and M-CM conditions comparing protrusion length (P = 0.012, n = 3), and cord count (P = 0.007, n = 3), in CG and MG conditions comparing protrusion thickness (P = 0.0000027, n = 3), number of protrusion (P = NS) and max branch length/mean branch length (P = 0.010, n = 3). Plotted is mean ± SD. Significance defined as P ≤ 0.05. Q) Phase contrast images on Day 7 of HepG2 spheroids in MG exposed to M-CM alone, M-CM with A83-01 (10 nm) and A83-01 (20 nm). R) Bar graph analysis comparing D7 M-CM and MCM + A83-01 (20 nm) (P = 0.047, n=3). Plotted is mean ± SD. Significance defined as P ≤ 0.05.

**Figure 5.**
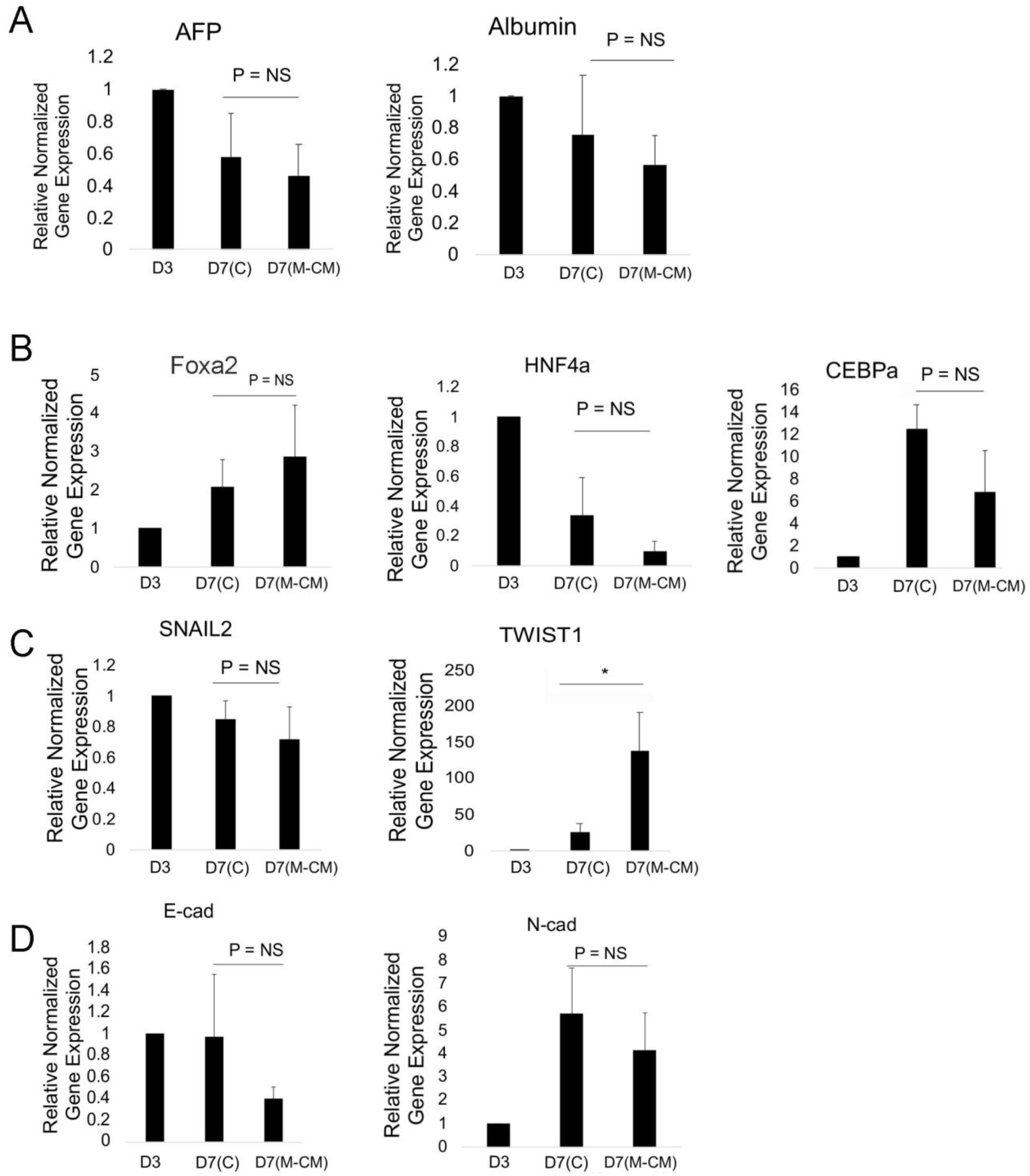
Gene expression analysis of HepG2 spheroids in MG droplets in M-CM. qRT-PCR analysis of HepG2 spheroids on day 3, and after culture in MG droplets with M-CM until Day 7. A) Markers associated with liver differentiation, including alpha-fetoprotein (AFP) (P = NS, n =3 for Day 7 Control, n=11 for D7 M-CM), albumin (Alb) (P = NS, n=3 for Day 7 Control, n = 9 for D7 M-CM) are shown. Plotted is mean ± SD. Significance defined as P ≤ 0.05. B) Transcription factors in liver development and maturation including Foxa2 (P = NS (P = 0.061), n = 3 for Day 7 Control, n = 5 for Day 7 M-CM), HNF4a (P = NS (P = 0.25), n = 3 for Day 7 Control, n = 10 for D7 M-CM), CEBPa (P = NS, (P = 0.103), n = 3 for Day 7 Control, n = 3 for Day 7 M-CM) are shown. Plotted is mean ± SD. Significance defined as P ≤ 0.05. C) Transcription factors associated with epithelial to mesenchymal transition (EMT) including SNAIL 2 (P = NS, n =3 for Day 7 control, n = 3 for D7 M-CM), and TWIST1 (P = 0.026, n =3 for Day 7 control, n = 4 for D7 M-CM) were tested. Plotted is mean ± SD. Significance defined as P ≤ 0.05. D) Cadherin expression associated with EMT, E-cadherin (E-Cad) (P = NS, (P = 0.14), n = 3 for Day 7 Control, n = 4 for D7 M-CM) and N-cadherin (N-Cad) expression (P = NS, n =4 for Day 7 Control, n = 4 for Day 7 M-CM). Plotted is mean ± SD. Significance defined as P ≤ 0.05.

**Figure 6.**
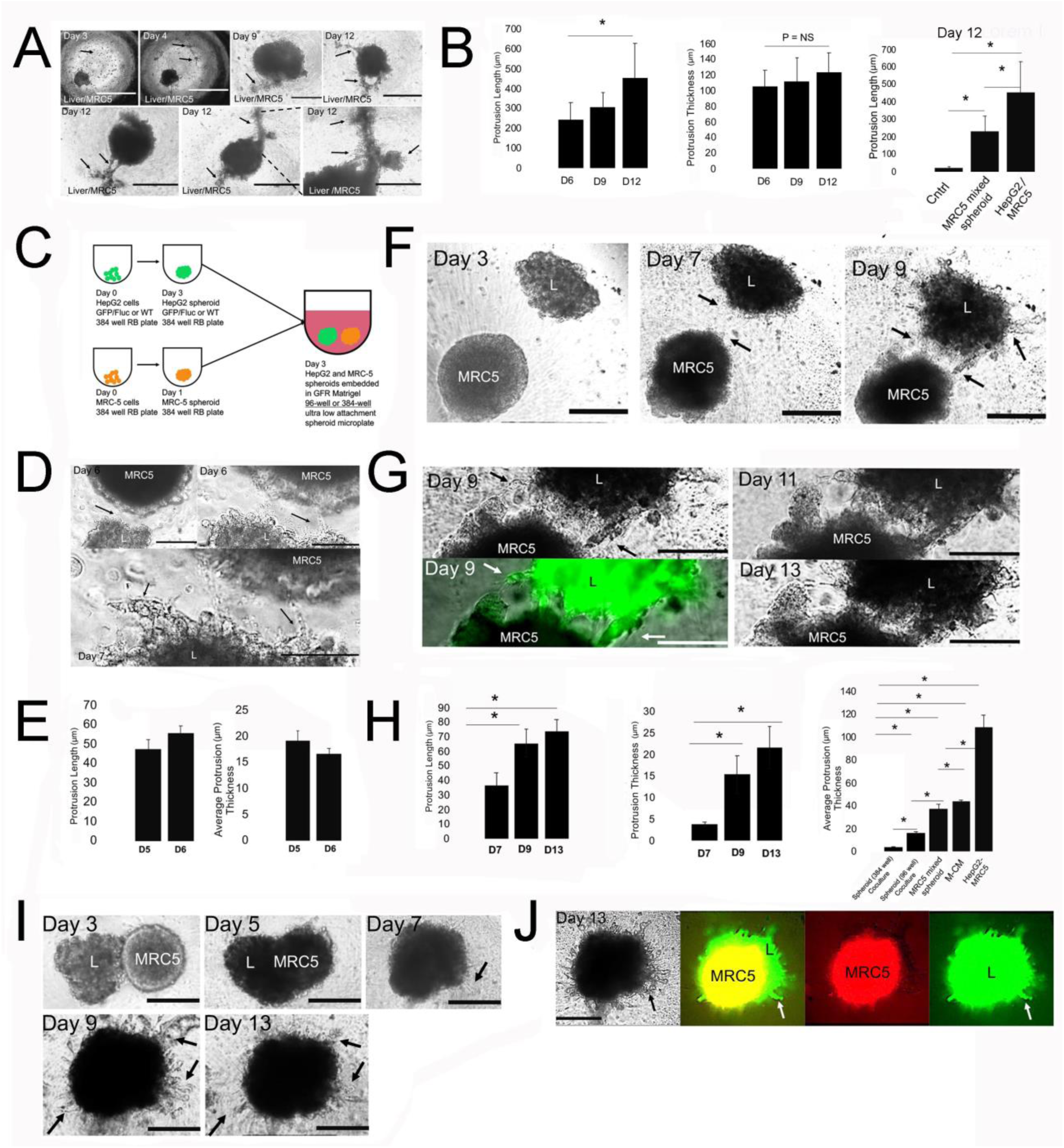
Engineering fibroblast density and format for increased 3D collective cell migration. A) Phase contrast microscopy images of HepG2 spheroids cultured in MG bearing high density (30,000 cells) of MRC-5 cells. Top row, left to right- Day 3, 4, 9, 12. Day 3- MRC-5 cells initially after seeding (arrow). Bar = 1000 µm. Day 4- MRC-5 cells spreading and interconnecting (arrows), Bar = 1000 µm. Day 9- Thick hepatic cord (arrows), Bar = 400 µm. Day 12- multiple, thick hepatic cords, Bar = 400 µm. Bottom row, left to right- Each image in a separate experimental replicate on Day 12 demonstrating thick hepatic cord formation (arrows), Bar = 400 µm. B) Bar graph analysis of HepG2 spheroids cultured in MG bearing high density (30,000 cells) of MRC-5 cells. Left to right, Left- Protrusion length on Days 6, 9, 12 shown, Day 9 and Day 12 compared, P = 0.05, n = 3. Middle- Protrusion thickness on Days 6, 9, 12 shown, Day 9 and Day 12 compared, P = NS, n = 3. Right -Bar graph analysis comparing Protrusion length between Control (HepG2 spheroid in MG), HepG2 and MRC-5 mixed spheroid, and HepG2 with high density MRC-5 in the MG. Control (n = 3) compared to HepG2 with MRC-5 in MG (n = 3), P = 0.00053, and to MRC-5 mixed spheroid (n = 3), P = 0.00085. MRC-5 mixed spheroid (n = 3) compared to HepG2 with MRC-5 in MG (n = 3), P = 0.0014. Plotted is mean ± SD. Significance defined as P ≤ 0.05. C) Schematic for modeling the liver diverticulum with co-spheroid culture. HepG2 spheroids and MRC-5 5 spheroids are cultured in 384-well nonadherent plates for days, and are either transferred together into a single well in a 96-well or 384-well and embedded in 50µL MG for several days of culture. D) Phase contrast images of co-spheroid culture in a 96-well plate. Top Left- Interface of liver spheroid (L) and MRC-5 spheroid (MRC-5) on Day 6. Arrow shows interacting structures, Bar = 200 µm. Top Right- high magnification view on Day 6. Arrow shows interacting structures, Bar = 100 µm. Bottom- Day 7 cultures. Arrows show migrating strands and interacting structures, Bar = 100 µm. E) Bar graph on days 5 and 6 demonstrating protrusion length and protrusion thickness. Plotted is Mean ± SD. Significance defined as P ≤ 0.05. F) Phase contrast images of co-spheroid culture in a 384-well plate for Days 3-9. “L” is the liver (HepG2) spheroid. Arrows demonstrate migrating hepatic cords between spheroids, Bar = 200 µm. G) Same study as in F) except Day 9- Day 13 phase contrast and fluorescent images shown. Left- High magnification of phase contrast (top) and corresponding fluorescent (HepG2-GFP) images of green anchoring strands (arrows). Arrow demonstrates migrating strands. Right-Phase contrast images of Day 11 (top) and Day 13 (bottom) images shown. Arrows demonstrate hepatic cords anchored to MRC-5 spheroid, Bar = 100 µm. H) Bar graph analysis of spheroid co-culture. Left- Protrusion length on Days 7, 9, and 11. Comparison of Day 7 and Day 9 (P = 0.00063, n =3), and Day 7 and Day 13 (P = 0.00011, n = 3) is shown. Middle- Protrusion thickness on Day 7, 9, and 13. Comparison of Day 7 and Day 9 (P = 0.004, n =3), and Day 7 and Day 13 (P = 0.0019, n = 3) is shown. Right- Comparison of protrusion thickness across all the culture systems developed on Day 6. Spheroid 384-well co-culture (Day 6, n = 3) compared to HepG2 with MRC-5 in MG (Day 6, P = 0.00357, n = 3), to M-CM (Day 6, P = 0.00007, n = 3) to MRC-5 mixed spheroid (Day 6, n = 3), P =0.00695, n = 3) and to Spheroid 96 well co-culture (Day 6, P = 0.00047, n = 3). We also compared Spheroid 96-well co-culture (Day 6, n = 3) to HepG2 with MRC-5 in MG (Day 6, P = 0.00046, n = 3), to M-CM (Day 6, P = 0.00059, n = 3), and to MRC-5 mixed spheroid (Day 6, P = 0.01868, n = 3). Plotted is mean ± SD. Significance defined as P ≤ 0.05. I) Phase contrast image of co-spheroid liver and MRC-5 spheroid on Days 3, 5, 7, 9, and 13. Day 3-5- spheroids interact, Day 7- spheroid fusion and migration at spheroid edge, Day 9- arrows demonstrate protrusions, Day 13- arrows demonstrate protrusions, Bar = 200 µm. J) Phase contrast and fluorescent images of fused HepG2 and MRC-5 spheroid. Left- Phase contrast (Day 13) image with arrows demonstrating protrusions. Double fluorescent (Green- HepG2-GFP, Red-MRC-5, arrows showing HepG2 migration), Red only (MRC-5), and Green- (“L”-Liver) with arrow showing HepG2 migration, Bar = 200 µm.

### Effects of 3D migration on gene expression

We performed gene expression analysis of liver spheroids in control and M-CM conditions on day 7 normalized to day 3 (liver spheroid). We analyzed genes associated with differentiation and migration. In the presence of M-CM, on day 7, we observed no significant changes in albumin and AFP expression (Fig 5A), while for key transcription factors, we observed a trending increase in Foxa2, a trending decrease in HNF4a (global transcriptional activator), and a significant decrease in CEBPa (metabolic functions) (Fig 5B). In terms of epithelial to mesenchymal (EMT) transcription factors, we observed no change in SNAIL2 expression, and significant upregulation of Twist1 (Fig 5C). We also observed strong trends downward for E-Cadherin expression, consistent with cell migration, as well as N-Cadherin expression (Fig 5D).

### Engineering fibroblast density and format for increased 3D collective cell migration

Although we consistently observe liver 3D CCM with MRC-5-conditioned medium, we wished to improve the *in vitro* modeling by again addressing how cellularity within the STM could affect cell migration and induce 3D liver CCM. We first increased the MRC-5 cellular density by 50% in the LD model (Fig. 1M). At these higher densities, we found that small islands of MRC-5 fibroblasts formed within the MG, and in response, thick liver spheroid-derived cords protruded from the spheroid to these fibroblasts islands to form thick strands containing both cell types (Fig. 6A). These cords formed by day 9 and thickened to an average of 115 microns by day 12 (Fig. 6B) and demonstrated significantly longer strands then the mixed spheroids and control (no MRC-5). We also observed that the liver spheroid exhibited radial migrating strands away from these thick strands at the edge of spheroids (Fig. 6A), but new strands did not form after day 9. To further increase cellular density of the MRC-5 fibroblasts and obtain a system in which migration occurred faster, we changed the experimental system to co-spheroid culture in which spheroids interact in MG (Fig. 6C). Recent studies demonstrate morphogenesis and collective cell migration through self-organization of multiple spheroids, but these were not submerged in a gel (Koike, Iwasawa et al. 2019). Surprisingly, we found that when spheroids were placed within 150 microns from each other in a 96-well plate, we observed filopodia extension with concomitant cell migration by day 6 (Fig. 6D). In this system we observed a trending increase in protrusion length and a trending decrease in protrusion thickness. (Fig. 6E). To improve spheroid interactions, we switched from a 96-well to a 384-well format. In this cultivation system, the liver spheroid is transferred from the well and there is a minor loss of circularity (Fig. 6F**, “L” labeled spheroid**). Here, we observed multiple thin strands/cords that form on day 9 (Fig. 6F). They originate from the liver spheroid and then they move towards the spheroid and anchor to the MRC-5 spheroid (Fig. 6G). This is highly reminiscent of hepatic cords that form during liver organ development (Compare Fig. 6F-G with Fig. 1A). We observed significant increase in protrusion length Fig. 6H**, left)** and protrusion thickness over time (Fig. 6H**, middle)**. When comparing all the systems we developed here for protrusion thickness, we observed statistically significant differences between the co-spheroid culture systems, and all other systems developed, with the co-spheroid culture demonstrating the thinnest strands (Fig. 6H**, right)**. In this co-spheroid culture system with a 384-well plate instead of a 96-well plate, we observed novel findings. From day 3 to day 7, we observed fusion of the two spheroids (Fig. 6I). This was followed by formation of 3D CCM of radial cell strands from days 7 to day 13 (Fig. 6I). These strands were maximal at day 9, but retracted by day 13. We performed fluorescent labeling and imaging of spheroids to understand spheroid interactions between liver spheroid (green) and MRC-5 spheroid (red). Interestingly, we found that the liver spheroid engulfs the MRC spheroid (Fig. 6J), from day 3 to day 7, using fluorescent imaging, which requires liver cells migrating interstitially over MRC-5 spheroids cells. This engulfing process occurs without an overt change in size, suggesting that cellular packing density is very tight within this spheroid. The two-color imaging enables us to identify that the cellular projections from the spheroid are primarily of hepatic origin (Fig. 6J). Minimal red fluorescent strands are seen, indicating that nearly all radial migrating strands are from the liver spheroid. Taken together, our data demonstrate that modeling the STM at high density results in significant morphogenetic events, such thick cord formation, thin cord formation, interstitial and anchoring migration, and spheroid fusion.

## DISCUSSION

Liver 3D CCM is critical in several scenarios, including: 1) hepatic cord formation during hepatic endoderm migration, 2) early fetal hepatocyte interstitial migration during liver expansion, 3) hepatocyte migration during liver repopulation, 4) spread and metastasis in hepatocellular carcinoma, and 5) other pathologic processes, like bridging fibrosis. We hypothesized that modeling the liver diverticulum can lead to improved 3D CCM modeling. Here, we employ well studied HepG2 cell lines which demonstrate an immature phenotype, and engineer both spheroid composition and mesenchyme composition to determine or highlight varying modes of 3D CCM, and enable improved visualization and analysis of migration. We varied both the mesenchyme and the spheroid properties to determine several novel cultivation systems in which 3D collective cell migration occurs. We developed, characterized, and in some cases identified molecular mechanisms for 3D CCM. The cultivation systems reported here include: 1) spheroids cultivated in hydrogels and mesenchymal cells, 2) mixed spheroids containing liver and mesenchymal cells, 3) liver spheroid migration with mesenchyme-conditioned medium, and 4) spheroid co-cultivation of liver and mesenchyme-derived spheroids. These systems further our understanding of paracrine factors and other physical means of cell-cell communication, thereby resulting in 3D CCM. Further, these systems enable modeling of various stages/forms of 3D CCM, which will further our molecular and cellular understanding of liver cell migration, and improve treatments for HCC metastasis and liver cell therapy.

A key finding is that M-CM initiates 3D liver migration in both MG and Collagen hydrogel systems. The MG studies demonstrates a model for branching morphogenesis, while the Collagen studies demonstrate linear and elongated protrusive growth. Our data also demonstrates that DAPI (nuclear) staining occurs in these protrusions, and that Ki-67 staining (proliferation) is associated with some of the protrusions. This data is consistent with study of HCC cell lines, in which M-CM initiates 2D migration of HCC cells cultured in monolayer, including changes in morphology, migration, and invasion, an increase in laminins, integrins, activated focal adhesion kinase and other downstream molecules, redistribution of cytoskeletal proteins, and changes in cell cycle (Ding, Chen et al. 2015). These published changes in migration can be classified as a non-classical EMT, and are tissue-specific and cell line-specific. For example, in pancreatic cancer cell lines, M-CM has been shown to inhibit cell migration, invasion, promote a more immunosuppressive phenotype, and coordinately increase cell polarity markers (Ding, Lu et al. 2018). The studies mentioned above both mention cell line-specific effects, and gene expression results that don’t correlate with protein expression, suggesting underlying complexity and protein redistribution as a mechanism. Interestingly, from a historical perspective, analysis of secreted factors of MRC-5 fibroblasts and other mesenchymal cells led to the isolation of HGF as a factor that independently causes cell migration (Stoker and Perryman 1985);(Schor, Schor et al. 1988)suggesting that HGF is a causative factor, but these studies do not identify a particular mechanism by which the M-CM functions in 3D migration. HGF has been shown to promote migration of murine oval cells, or hepatic stem/progenitor cells (Suarez-Causado, Caballero-Diaz et al. 2015), as well as Huh-7 and HepG2 cells (Meng, Wei et al. 2015) using traditional *in vitro* assays. Since TGFß pathway is implicated in hepatic cord formation (Rossi, Dunn et al. 2001) and HCC migration (Fransvea, Angelotti et al. 2008), and has shown to be involved in cross-talk with HGF pathway (Rossi, Dunn et al. 2001), we evaluated TGFß pathway inhibitors. We indeed identified that large effects in migration can be attributed to TGFß-mediated 3D CCM (Fig. 2D). The addition of A83-01, a broad inhibitor which inhibits TGFß, Activin and Nodal signaling pathway, by inhibiting ALK4, ALK5, ALK7 receptors and preventing SMAD2/3 phosphorylation, resulted in a dramatic decrease in migration in a dose-dependent fashion (Fig. 4Q). Although we observed increased migration by employing HepG2 cells, it remains to be seen if early and late stage human hepatoblasts, and human hepatocytes, respond in a similar way. Further, analysis employing systems biology techniques of the secretome and exosome of M-CM that may find the mechanism by which MRC-5 functions. Human MRC-5 lung fibroblast cells were isolated at 14 weeks which corresponds to approximately E18 in the mouse, which suggests that fetal-like mesenchyme may have potent effects on 3D CCM. In terms of STM modeling, human stem cells that have been engineered to mesenchymal cells that mimic the STM are in its earlier stages, and employing them in *in vitro* models of liver bud has commenced (Takebe, Sekine et al. 2017). Studies continue to elucidate key molecules underlying the STM phenotype (Duenas, Exposito et al. 2019) which suggests that improved *in vitro* modeling of the mesenchymal cells can be accomplished using human stem cells (Coll, Perea et al. 2018).

The factors that promote liver co-migration of liver and mesenchymal cells are poorly understood. The concept of co-migration is critical because co-migration of hepatic endoderm, STM cells, and endothelial cells occurs in the liver bud, after the liver diverticulum stage (Cascio and Zaret 1991). We determined the conditions under which co-migration occurs using a mixed spheroid system containing liver cells and either HFF or MRC-5. Mixed liver spheroids have been previously used to improve viability of spheroids, and enhanced cancer invasiveness, as demonstrated by changes in gene expression and enhanced drug resistance (Jung, Kang et al. 2017). Further, mixed systems have been shown to model the liver bud, which is the stage immediately after the liver diverticulum stage (Takebe, Sekine et al. 2013) (Fig. 1A). The mixed spheroids in MG resulted in co-migration with HFF (Fig. 2F). When we employed stiffer matrix, like collagen and fibrin, we observed extremely long, thin protrusions that were submicron and hair-like. We used cell labeling to determine that these long thin protrusions are fibroblasts and separately liver cells (Fig. 2H, Supp. Fig. 4), and therefore not co-migration. This suggests that co-migration requires either soft hydrogels or components within the MG. Further, in our co-migration models, we don’t observe branching morphogenesis, as we do with M-CM studies. It remains to be seen if co-migration together with branching morphogenesis can occur simultaneously. Our data suggests that TGFß1increases the number of protrusions and the overall area of growth due to co-migration, but we don’t observe branching in any of these cases of co-migration. The factors that influence liver/mesenchymal co-migration are not known. However, in studies of squamous cell carcinoma, it was shown that Rho-ROCK activated fibroblasts physically lead collective cell migration by secreting proteases (MMP) and generating tracks within the collagen matrix in which groups of cancer cells can follow (Gaggioli, Hooper et al. 2007). We feel by improved modeling of various aspects of morphogenesis, then we will be able to better tease out the multiple biomechanical and protein translational changes that result in morphogenesis and/or migration (Okuda, Takata et al. 2018), and use this to improve liver cell therapy and cancer therapies.

An interesting form of the mixed spheroid condition is when the liver spheroid fuses with the MRC-5 spheroid (Fig. 6I-J). In the absence of spheroid fusion, we observed thin strands originating in the liver spheroid and interacting with the MRC-5 spheroid, very similar to what occurs within the liver diverticulum. These cell strands or protrusions do not contain branches, and anchor to the MRC-5 spheroid indicating morphogenetic processes and interstitial migration. In some cases, these self-assembled spheroids undergo fusion, which we term self-driven morphogenesis (Sasai 2013). This is highly reminiscent of fused organoids that have received increased attention (Bagley, Reumann et al. 2017);(Xiang, Tanaka et al. 2017);(Xiang, Tanaka et al. 2019);(Mansour, Goncalves et al. 2018) for stem cell biology and bioprinting (engineering) applications. Importantly, employing spheroid fusion of stem-cell derived foregut and midgut organoids, hepatic, biliary and pancreatic organ tissue arises (Koike, Iwasawa et al. 2019), and the authors point out that migration of hepatic endoderm progenitors did not occur. This points to the need for liver 3D CCM in the setting of spheroid fusion. Here, we demonstrate clear evidence of hepatic cells engulfing mesenchymal (MRC-5) spheroids, which likely requires interstitial, 3D CCM of liver cells migrating above MRC-5 fibroblasts. A physiological example of a similar process occurring, in which an epithelial population engulfs a mesenchymal population, is during the process of hair follicle development (Millar 2002), specifically when the inner root sheath is formed. Similarly, it has been shown that spheroids of retinal progenitor cells (epithelial cells) can envelope spheroids of limbal MSC in culture (Kosheleva, Efremov et al. 2020), consistent with our MRC-5 fibroblasts remaining inside. In terms of mechanism of spheroid fusion, it was previously found that that tissue spheroids that were exposed to TGFß favored fusion with non-exposed spheroids, which was consistent with our data that MRC-5 cells may secrete TGFß molecules (Hajdu, Mironov et al. 2010). The roles for extracellular MG, signaling pathways, and cell and tissue mechanics still needs to be determined. It remains unknown whether this process occurs during liver development, regeneration, or disease. However, two morphogenetic structures which are important to speculate about are: 1) regenerative nodules that occur during chronic liver disease, and 2) bridging fibrosis which leads to advanced stage fibrosis and cirrhosis. We speculate that the 3D models here could be further developed to model these more complex processes. Since the liver is a metabolically active cell, it would be interesting to determine the energy requirements for liver cell migration, and the significance of liver disease, as suggested by recent studies (Zanotelli, Rahman-Zaman et al. 2019). Importantly, the fused spheroid system we have developed results in liver cell migration, but not co-migration. These are duct like protrusions that are uniquely liver only, that do not branch, and whether the mechanism is chemical (TGF) or biomechanical (stiff mesenchymal core) is unclear.

There were several limitations to our study. We observed migration followed by branching morphogenesis in the presence of M-CM. A major question is whether nuclei also migrate in the structures. Our data of DAPI staining strongly suggests that at least some of these structures have nuclei (Supp. Fig 8). Further analysis of nuclear staining at high resolution is needed to confirm the nature of these branching structures. From a methods point of view, it was challenge to recovering spheroids from miniature plates like 96 and 384-well plates. This resulted at times in a slight loss of sphericity, and we attempted several approaches are still developing approaches to transfer spheroids from one system to another. Our studies of extracellular matrix effects tested a few types of matrix, but future studies will analyze the effects of extracellular matrix more vigorously, including studies of fibrillar vs cross-linked collagen. Our MG droplet system lasts about 2 weeks, but then the matrix degrades significantly, and therefore new methods are needed to improve this system for long term culture. Finally, as our systems were in well plates and tissue culture plastic-based plastic/wells, it was challenging to perform high magnification of analysis, which will be performed on the future on glass for improved imaging, or with tissue clearing protocols followed by light-sheet or multiphoton microscopy. Nonetheless, we present several new systems which result in various forms of 3D CCM. We not only observe varying thickness and length of protrusions that can be further dissected mechanistically, we also demonstrate co-migration, branching morphogenesis, and interstitial migration, all of which are valuable for scientists modeling liver development, liver cell therapy, or liver diseases.

## Competing Interests

The authors hereby state no competing interest involved with the ideation, writing, or revision of this manuscript

## Acknowledgements

NP was supported by the UB CBE startup funds, the New York State Stem Cell Science C024316 and the Stem cells in regenerative medicine (ScIRM) center. OO was supported by the Western NY Prosperity Fellowship.

## AUTHOR CONTRIBUTIONS

OO: Obtained data, analyzed data, wrote and approved manuscript

OY: Obtained data, analyzed data, wrote and approved manuscript

CO: Obtained data, analyzed data, approved manuscript

AK: Obtained data, analyzed data, approved manuscript

LW: Obtained data, analyzed data, approved manuscript

CS: Obtained data, analyzed data, approved manuscript

SR: Obtained data, analyzed data, approved manuscript

NP: Conceptualized, acquired funding, investigated, supervised, wrote, edited manuscript, and approved manuscript.

## Supplemental Figure Legends

**Supplementary Figure 1.**
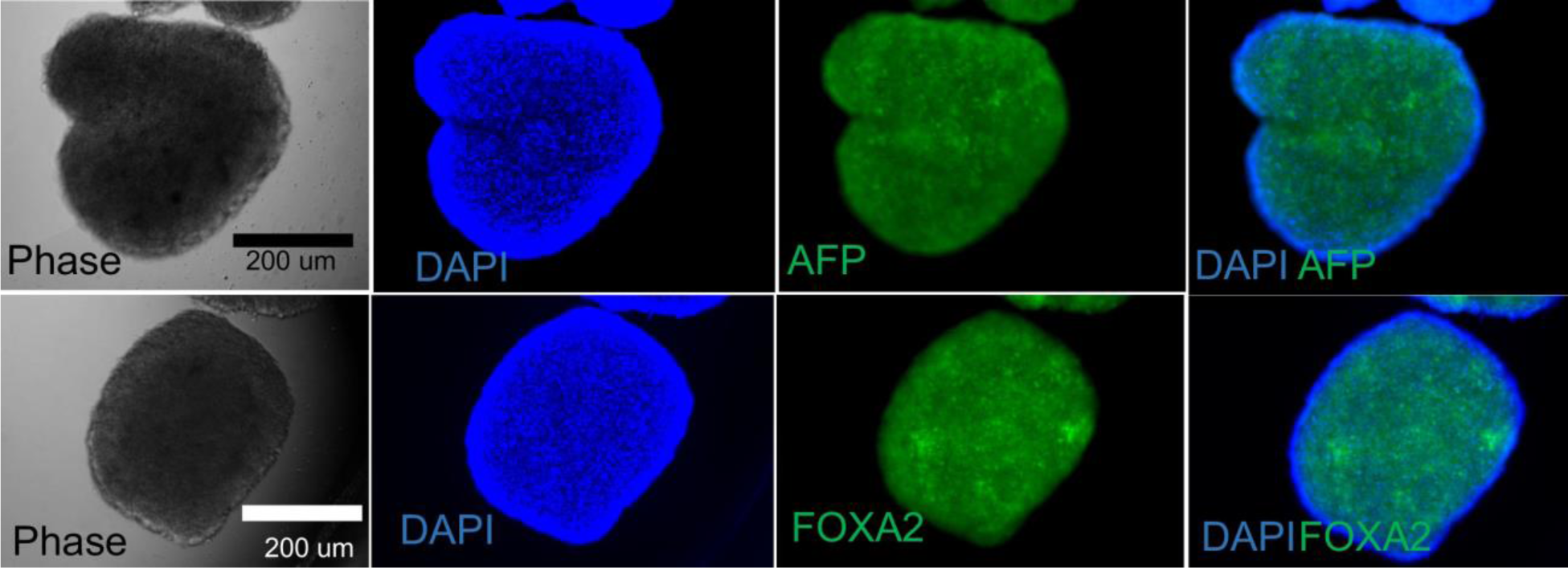
HepG2 spheroids in suspension culture were collected and immunolabeled in suspension. Briefly, spheroids were harvested and fixed with 4% paraformaldehyde (PFA) at room temperature, permeabilized with 1% Triton X-100 and then incubated overnight with either alpha-fetoprotein (AFP) or FOXA2 monoclonal primary antibody. These markers are specific to liver lineage commitment. HepG2 spheroids were observed to positively express FOXA2 and AFP.

**Supplementary Figure 2.**
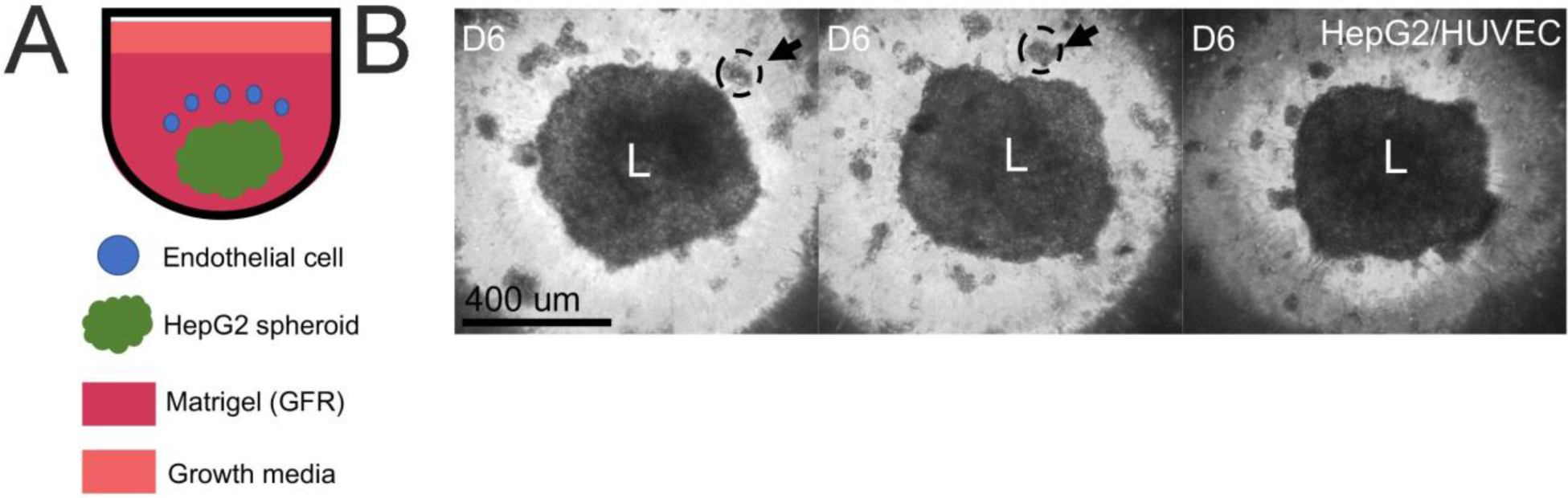
HepG2 spheroids (L) were initially formed in suspension culture using 384 well round bottom ultra-low attachment plates (1000 cells per well). On Day 3 spheroids were then embedded in Matrigel (GFR) containing 20,000 HUVEC cells. Additional growth media to support HepG2 and endothelial cell proliferation, DMEM: EGM in a 1:1 ratio, was used to culture the embedded spheroids. By Day 6 no significant outward migration of HepG2 cells was observed into the ECM. Aggregates of endothelial cells were observed in the Matrigel as clusters (black arrow) that showed no interaction with the HepG2 spheroid.

**Supplementary Figure 3.**
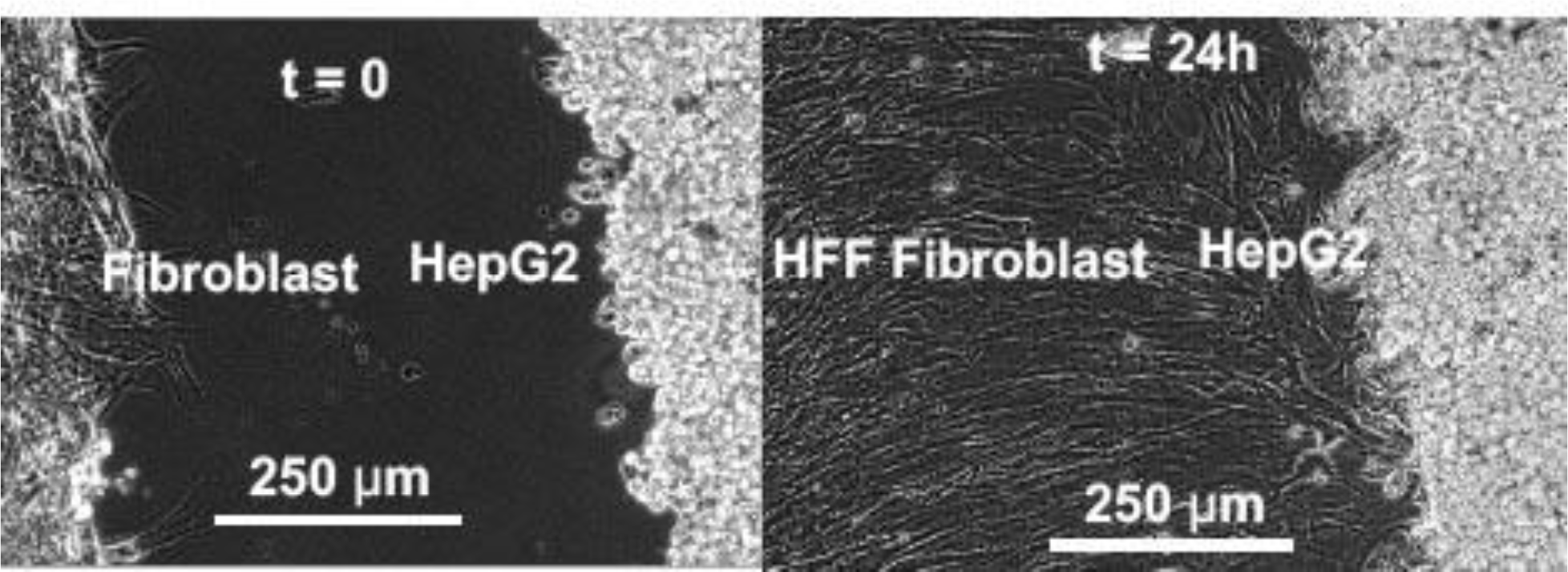
A barrier migration assay to establish the effect of human foreskin fibroblasts (HFFs) on HepG2 collective cell migration. Briefly, an ibidi™ 2 well insert was used to create a barrier between HFFs and HepG2 cells seeded 500 µm apart. After overnight incubation to allow for cell attachment, the insert was removed allow for migration between the cell reservoirs. There is observed collective migration of HepG2 cells outwards in response to the presence of the HFFs present.

**Supplementary Figure 4.**
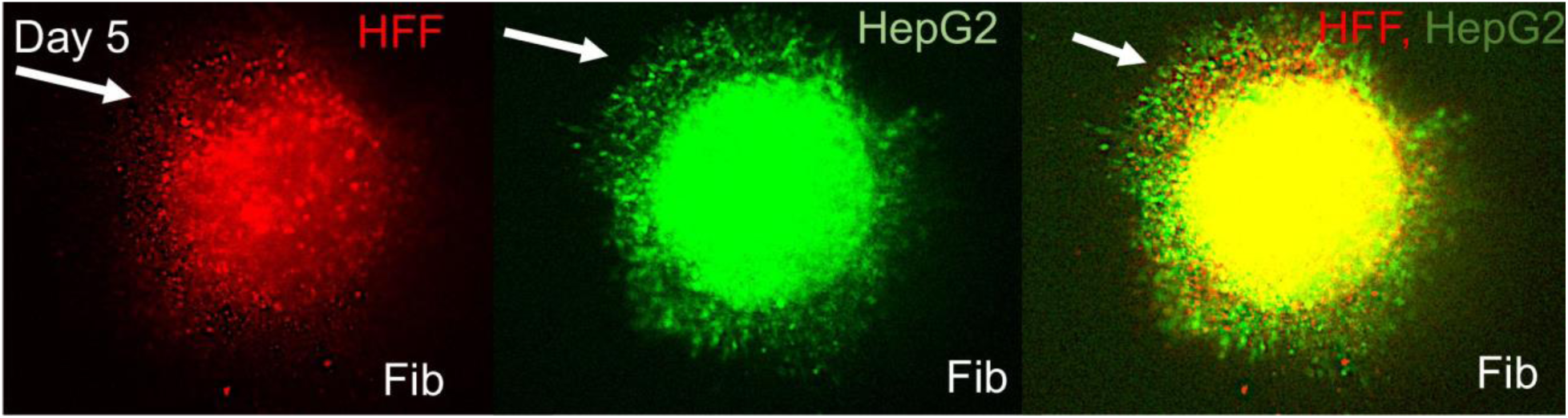
HepG2 cells and Human foreskin fibroblasts (HFFs) were initially separately dye labeled green and red respectively and then mixed in a 1:1 ratio before being seeded into 384 round bottom ultra-low attachment plates. Cells formed a compact spheroid that was subsequently embedded in a pre-made fibrin hydrogel. Fibrin hydrogel was made by polymerizing fibrinogen (3.25 mg/mL) with thrombin (12.5 U/mL) in a 4:1 ratio. There was enhanced co-migration of HepG2 cells and HFF cells outwards into the fibrin matrix.

**Supplementary Figure 5.**
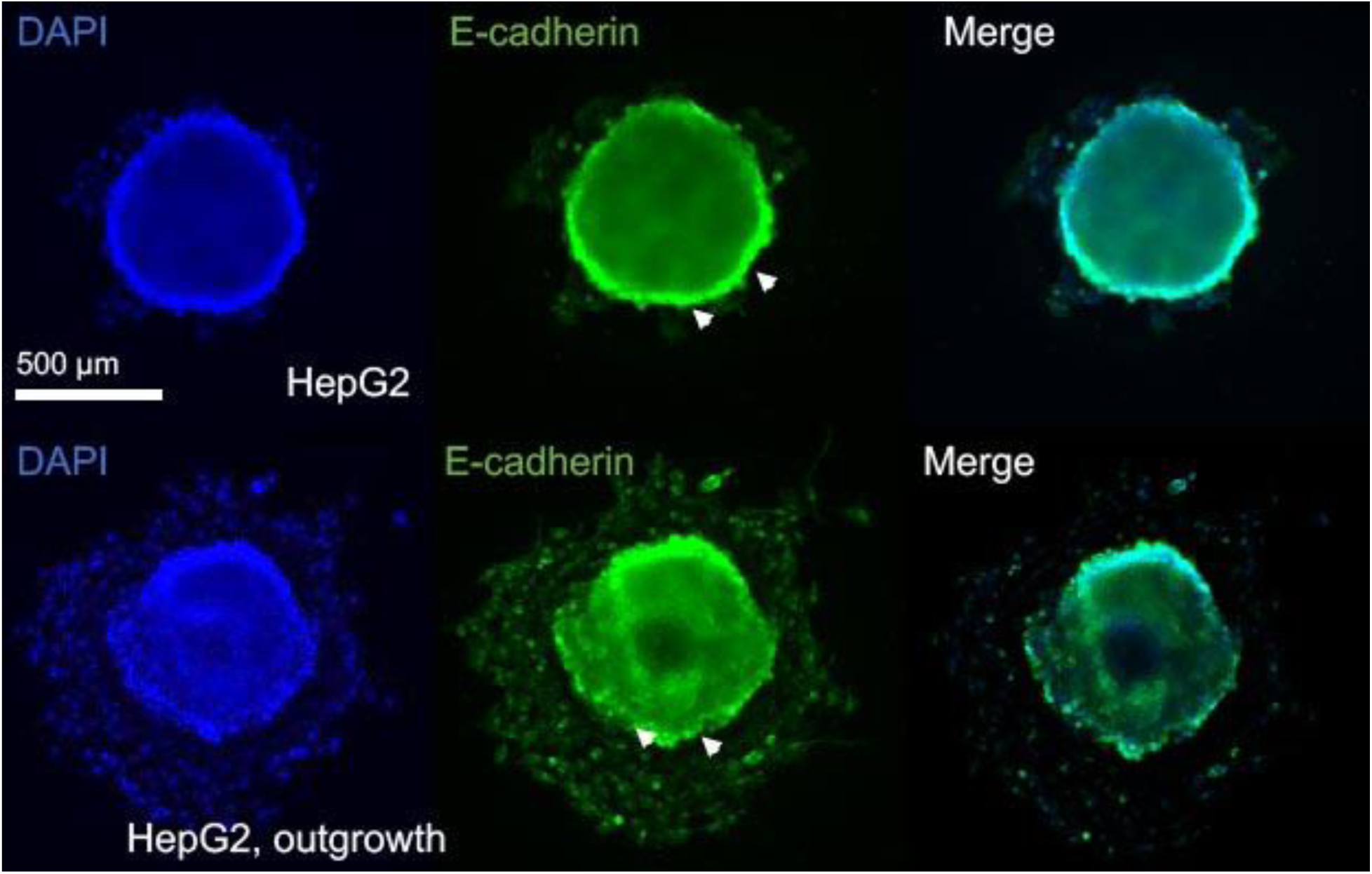
HepG2 spheroids seeded onto Matrigel coated plates in the presence of MRC-5 conditioned media (MCM) were observed to significantly migrate outwards as compared to control, serum containing DMEM. After 24 hours in culture, spheroids were fixed in 4% paraformaldehyde, permeabilized with 1% Triton X-100, and immunolabeled with a monoclonal antibody for E-cadherin. Spheroids cultured in MCM showed irregular localization of E-cadherin around the migrating edge of the spheroid, suggestive of collective cell movement outwards. Spheroids cultured in the control condition, exhibited strong e-cadherin expression along the spheroid edge.

**Supplementary Figure 6.**
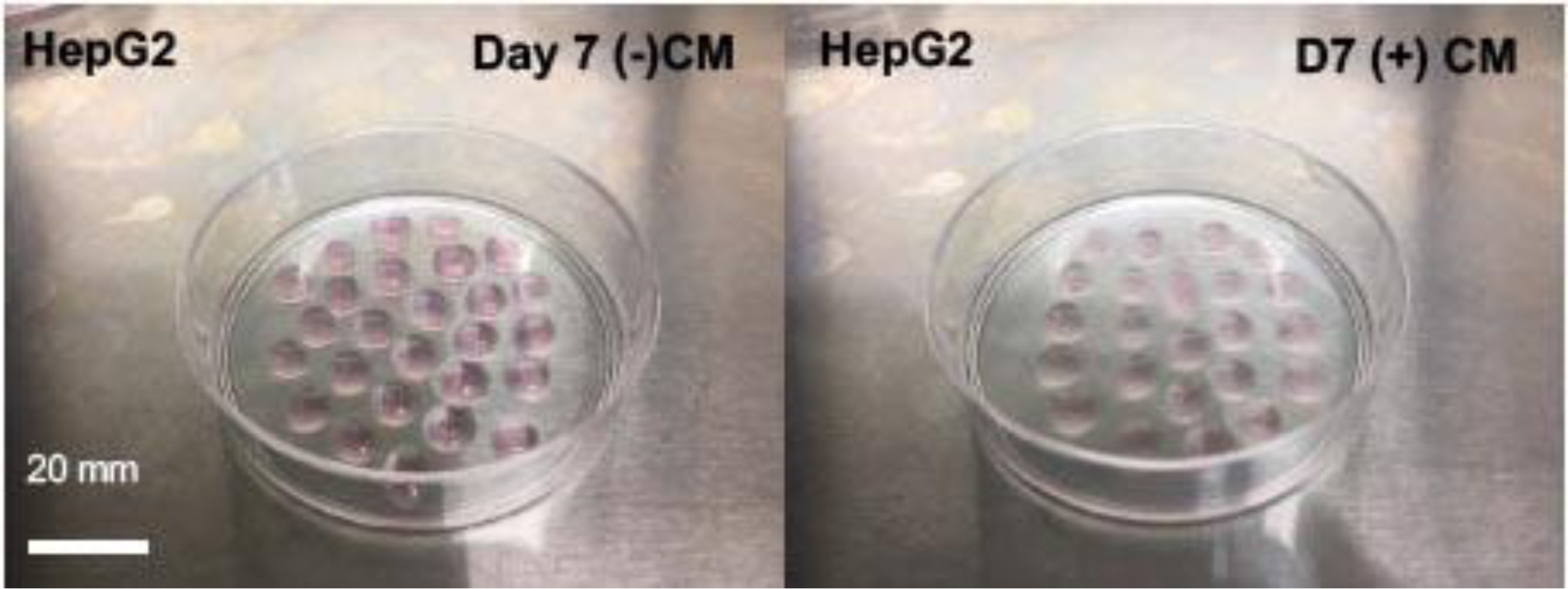
Images obtained of Day 7 HepG2 matrigel droplet culture systems in MRC-5 conditioned Media (MCM) and control experimental conditions (serum containing DMEM). The MCM condition is a lighter pink color than the control condition which is supportive evidence for the degradation of the matrigel by the migratory HepG2 cells in response to the presence of the MCM.

**Supplementary Figure 7.**
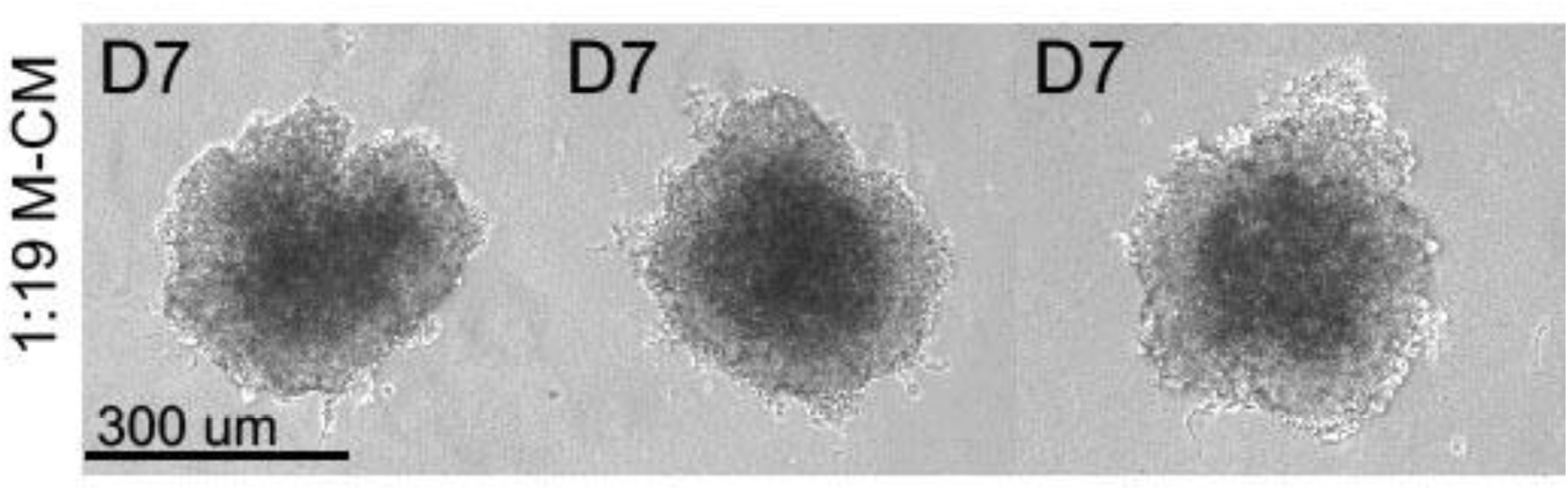
HepG2 spheroids cultured in 1:19 diluted MRC-5 conditioned media (M-CM) exhibit no matrix invasion. HepG2 spheroids formed in suspension culture using a 384 round bottom ultra-low attachment were harvested and seeded into Matrigel droplets on tissue culture treated 60 mm dishes. Matrigel droplets were then cultured in 1:19 M-CM for an additional 4 days. On Day 7 spheroids showed no significant outwards migration into the extracellular matrix (ECM). The M-CM media typically supports extensive HepG2 migration.

**Supplementary Figure 8.**
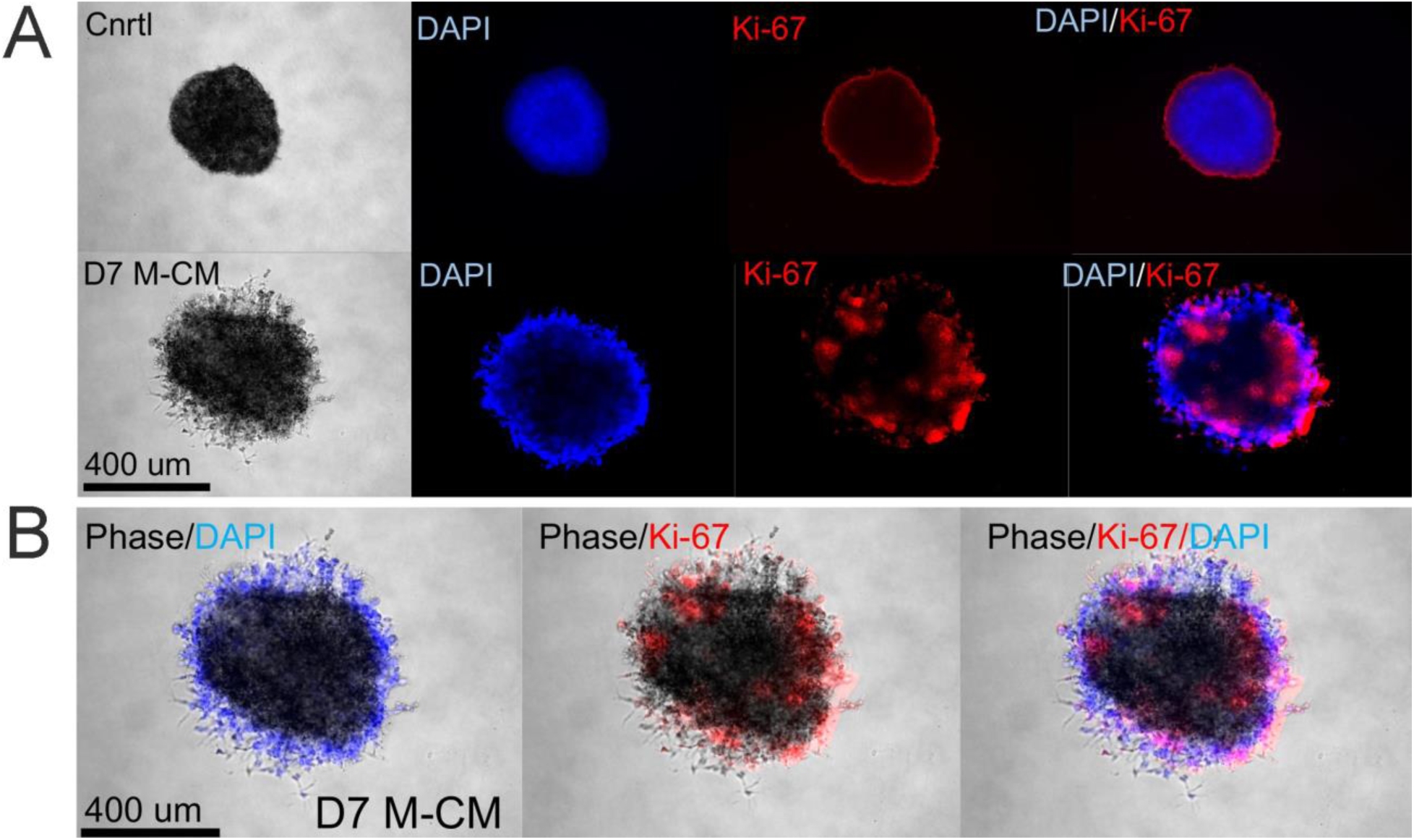
Ki-67 staining of HepG2 spheroids in Matrigel droplet culture. HepG2 spheroids formed in suspension culture using a 384 round bottom ultra-low attachment were harvested and seeded into Matrigel droplets on tissue. Culture treated 60 mm dishes. Matrigel droplets were then cultured in MRC-5 conditioned media (M-CM) for an additional 4 days. On Day 7 droplets were fixed with 4% paraformaldehyde (PFA) at room temperature, permeabilized with 1% Triton X-100 and stained using monoclonal primary antibody, Ki-67, a marker of proliferation. Spheroids treated with M-CM staining showed positive expression for Ki-67 localized to migrating cords as compared to control, droplets cultured in serum containing DMEM.

## Notes

### Competing Interest Statement

The authors have declared no competing interest.

